# Genome editing of phylogenetically distinct bacteria using portable retron-mediated recombineering

**DOI:** 10.1101/2025.06.16.660010

**Authors:** Alejandro González-Delgado, Laura Bonillo-Lopez, Milo S Johnson, Nastassia Knödlseder, Ching-Chung Ko, Yassir Lekbach, Jee-Hwan Oh, Hemaa Selvakumar, Michael C Wold, Zihan Yu, Virginia Aragón, Jeffrey A Gralnick, Marc Güell, Graham F Hatfull, Benjamin K Keitz, Britt Koskella, Vivek K Mutalik, Jan-Peter van Pijkeren, Seth L Shipman

## Abstract

Advanced genome editing technologies have enabled rapid and flexible rewriting of the *Escherichia coli* genome, benefiting fundamental biology and biomanufacturing. Unfortunately, some of the most useful technologies to advance genome editing in *E. coli* have not yet been ported into other bacterial species. For instance, the addition of bacterial retrons to the genome editing toolbox has increased the efficiency of recombineering in *E. coli* by enabling sustained, abundant production of ssDNA recombineering donors by reverse transcription that install flexible, precise edits in the prokaryotic chromosome. To extend the utility of this technology beyond *E. coli*, we surveyed the portability and versatility of retron-mediated recombineering across three different bacterial phyla (*Proteobacteria, Bacillota* and *Actinomycetota*) and a total of 15 different species. We found that retron recombineering is functional in all species tested, reaching editing efficiencies above 20% in six of them, above 40% in three of them, and above 90% in two of them. We also tested the extension of the recombitron architecture optimizations and strain backgrounds in a subset of hosts to additionally increase editing rates. The broad recombitron survey carried out in this study forms the basis for widespread use of retron-derived technologies through the whole Bacteria domain.

## INTRODUCTION

Bacterial communities play a central role in nearly every ecosystem on Earth. Genome editing technologies that modify bacteria can, therefore, help us understand and intervene in these communities to promote sustainable agriculture (Gordon et al., 2021), increase food security (Pixley et al., 2022), and improve human health (Li et al., 2020). In the last decades, multiple approaches have been developed to edit bacterial genomes including traditional homologous recombination and more recently CRISPR-Cas based tools (Arroyo-Olarte et al., 2021). Yet, despite substantial progress, precise, scarless edits of one to hundreds of base pairs in length remain difficult to achieve efficiently in many species, underscoring the necessity for alternative genome editing techniques specially in non-model bacteria (Vento et al., 2019).

One of the most powerful approaches to edit bacterial genomes is known as recombineering (Wannier et al., 2021). Recombineering was developed in *Escherichia coli* 25 years ago to allow precise gene knockouts and point mutations (Datsenko et al., 2000; Yu et al., 2000; Ellis et al., 2001). It relies on the activity of a single-stranded annealing protein (SSAP) together with a single-stranded binding protein (SSB) to integrate foreign ssDNA into the lagging strand of the host chromosome during replication (Ellis et al., 2001; Mosberg et al., 2010; Wannier et al., 2020; Filsinger et al., 2021). In addition to individual mutations, a high-throughput version of recombineering called MAGE (Multiplexed Automated Genome Editing) achieves directed mutagenesis of defined genomic loci by electroporation of libraries of synthetic recombineering oligonucleotides (Wang et al., 2009; Ronda et al., 2016).

Unfortunately, recombineering is not without limitations. Recombineering requires delivery of exogenous DNA, typically synthesized oligonucleotides, which constrains the efficiency and broader applicability of this technology beyond *E. coli* (Corts et al., 2022). In addition, the delivered DNA is rapidly degraded by host nucleases so multiple cycles of electroporation are required to reach useful genome editing efficiencies (Wannier et al., 2021; Isaacs et al., 2011).

In the last years, the addition of bacterial retrons to the recombineering machinery has increased efficiency and extended the overall approach. Retrons are bacterial tripartite immune systems composed of a reverse transcriptase (RT), a non-coding RNA (ncRNA) with two regions known as msr and msd, and an additional protein or RT-fused domain that acts as an effector for the defense response (Millman et al., 2020; Gao et al., 2020; Mestre et al., 2020). The retron RT specifically recognizes its ncRNA and reverse transcribes the msd component into ssDNA (Inouye et al., 1989; Lim and Maas, 1989). For bacterial defense, retrons work as toxin/antitoxin systems (Millman et al., 2020; Gao et al., 2020; Bobonis et al., 2022; Palka et al., 2022). In the absence of the viral attack the ssDNA can repress the action of the effector protein, but upon phage infection this repression is relieved and the active effector is able to kill or inhibit the growth of the infected bacteria, shielding the cell population from the invading agent (Bobonis et al., 2022; Carabias et al., 2024).

In the context of recombineering, the retron effector is omitted and the retron ncRNA is modified so that the msd encodes an editing donor. When supplied on a plasmid along with the recombineering SSAP and SSB, this leads to sustained production of multiple copies of a specific ssDNA donor *in vivo* which is used in place of an oligonucleotide donor by the rest of the recombineering machinery (**Fig 1a**; Farzadfard et al., 2014; Schubert et al., 2021; Lopez et al., 2021). The combination of the prior recombineering components (SSAP/SSB) with the retron components has been called a recombitron. Recombitrons have led to increased efficiency of recombineering in *E. coli* due to the sustained, abundant donor ssDNA. Additionally, they have enabled new applications that were not possible with standard recombineering including (i) molecular recording, in which the production of an editing donor is driven by a biological signal, (ii) genome-wide functional genetic variants, in which the presence of a recombineering donor in the retron ncRNA on a plasmid serves as a proxy for the edit in the strain, (iii) phage editing, in which a phage is edited while replicating through a host expressing the recombitron components, (iv) metabolic engineering, in which multiple edits are made to an individual genome from a multiplexed recombitron cassette, and (v) gene-specific continuous evolution, in which multiple variants of structural domain or small proteins could be created in combinations with error-prone polymerases (Farzadfard et al., 2014; Schubert et al., 2021; Fishman et al., 2023; Liu et al., 2023; Gonzalez-Delgado et al., 2024; Liu et al., 2024).

**Figure 1.**
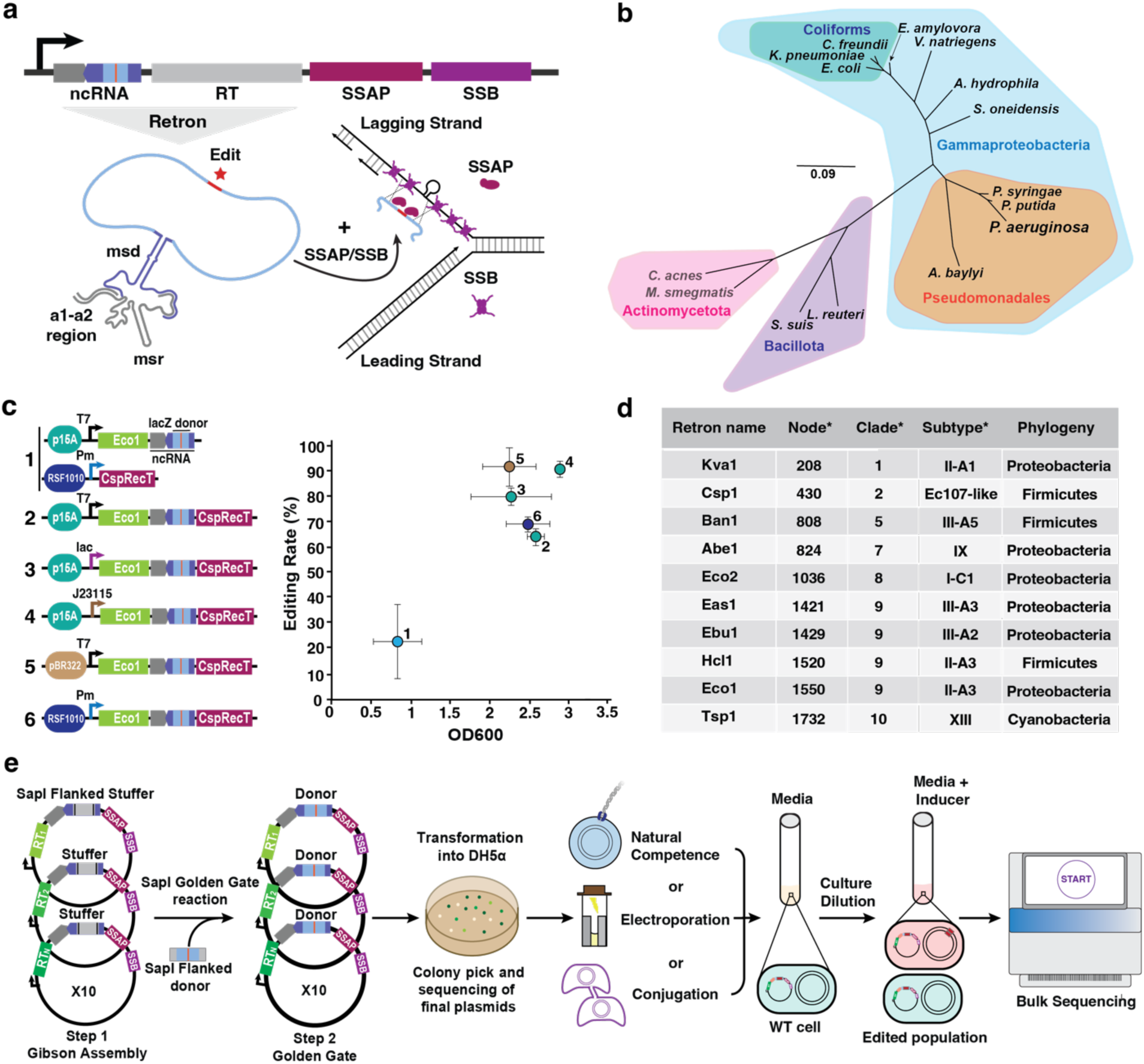
Design and construction of recombitrons for editing across the domain of Bacteria. **a**, Top: schematic of the recombitron operon with a donor encoded within the msd region of the ncRNA. Bottom: schematic of the retron-mediated recombineering process. Briefly, the retron RT produce multiple copies of the RT-DNA donor. SSAP and SSB proteins promote the binding of the RT-DNA donor to the lagging strand during bacterial replication installing the desired mutation. **b**, Unrooted phylogenetic tree of the bacterial species used in this work. The tree was constructed using a multiple sequencing alignment (MSA)of the 16S sequences of these species. The different colors represent the different bacterial phyla (Proteobacteria, Bacillota and Actinomycetota), class (Gammaproteobacteria), order (Pseudomonadales) or functional group (Coliforms). **c**, Left: Schematic of the operons used for optimizing recombitron architecture. Eco1 recombitron to make a deletion in the lacZ gene was used in all the configurations. Plasmid origin of replication and promoters are indicated in different colors. Right: quantification of precise editing rates of the lacZ locus and correlation with OD_600_ of the cultures after overnight growth. Editing data were quantified by Illumina sequencing. Error bars are ± standard deviation for three biological replicates. **d**, Table summarizing the relevant features of the retrons selected to perform the retron-based recombineering experiments. Information provided in the column with an asterisk was sourced from Mestre et al., 2020. **e**, Schematic of the workflow used to analyze the portability of recombitrons. Briefly, the set of 10 recombitrons was cloned in a host-specific plasmid using a Gibson Assembly approach. SapI flanked donors were ligated in the retron ncRNA of the 10 retrons in parallel employing a Golden Gate reaction. Multiple colonies were screened using Sanger sequencing to obtain the different recombitrons with the proper donor in a specific plasmid backbone. Plasmids were naturally introduced, electroporated, or conjugated into the final host. Single colonies are grown until saturation, diluted (with the proper inducer if required) and grown again until saturation. Precise editing rates are measured by Illumina sequencing.

Yet, despite its potential, retron-mediated recombineering has so far been confined to *E. coli*. Here, we explore the extension of retron recombineering across many species to identify new opportunities and challenges. To facilitate such a broad portability search, we used a multi-site design for this study. Recombitron parts were designed and constructed centrally, then shipped for testing by experts in each individual species at external sites, after which material (cell pellets or DNA) was shipped back to the central site for standardized analysis by sequencing. With this large coordinated collaboration between nine research labs, we screened recombitrons in a total of 15 bacterial species from 3 different phyla, including some in which recombineering technologies have never been tested. We found that recombitrons are functional in all species tested, reaching editing efficiencies above 20% in six of them, above 40% in three of them, and above 90% in two of them. We also tested the extension of recombitron architecture optimizations and strain backgrounds in a subset of hosts to additionally increase editing rates. The broad recombitron survey carried out in this study lays the groundwork for the use of retron-derived technologies through the whole Bacteria domain.

## RESULTS

### Design and construction of recombitrons for editing across the domain of Bacteria

To test the portability and versatility of retron-based recombineering, we selected a set of phylogenetically diverse species including Gram-negative, Gram-positive and *Mycobacteria* to perform editing experiments (**Fig 1b**; Supplementary Table 1). The different species were chosen based on their relevance as model organisms (e.g., *Escherichia coli*), their use as synthetic biology chassis (e.g., *Vibrio natriegens*, *Pseudomonas putida*), their relevance in industry (e.g., *Citrobacter freundii*, *Limosilactobacillus reuteri, Acinetobacter baylyi*), their potential to cause human disease (e.g., *Klebsiella pneumoniae, Pseudomonas aeruginosa, Aeromonas hydrophila Streptococcus suis*,), or plant pathogenicity (e.g., *Pseudomonas syringae*). An additional consideration was genetic access and the availability of previously tested molecular parts, like plasmids and promoters, that enable these experiments. Ultimately, a total of 15 bacterial species were included in the retron recombineering survey in this work. To facilitate such a broad portability search, we used a centrally coordinated, multi-site design (Supplementary Fig 1a).

Prior to editing beyond *E. coli*, we first standardized a recombitron operon architecture with the aim of achieving a single plasmid system and an origin of replication compatible with as many species as possible. Recent optimizations in plasmid architecture for retron editing have increased recombineering editing rates to nearly 100% in *E. coli* (Liu et al., 2023; Ni et al., 2024), which we leveraged for the design of our single plasmid recombitron systems. We first compared two plasmid recombitron designs used in prior work (López et al., 2021) to single plasmid designs each based on retron-Eco1 and the SSAP CspRecT for introducing a 10-base pair (bp) deletion in the *lacZ* gene in *E. coli.* We tested multiple origins of replication (p15A, pBR322, RSF1010), as well as inducible (T7, lac) and constitutive (RSF1010) promoters. All single plasmid designs outperformed the two plasmids system (**Fig 1c**). In addition to higher editing, cultures expressing the recombitron from a single plasmid reached a higher OD, indicating lower cellular burden of the single plasmid systems. These single plasmid recombitrons yielded editing rates up to 92%— the highest reported rate for a 10 bp deletion using retrons to date—with moderate differences in editing and growth when using different origins and promoters. Despite slightly lower editing rates, we chose to move forward with a single plasmid recombitron based on a broad host range RSF1010 origin of replication to facilitate porting into other species, and an inducible Pm promoter to avoid constitutive expression during cloning of subsequent editors.

To maximize the chance of successfully porting the system to other species, we chose to test 10 recombitron versions in each new species, based on 10 different retrons. Our research group recently completed an experimental census of retrons in which many systems were tested for their ability to produce DNA and support genome editing (Supplementary Fig 1B; Khan et al., 2024). From this census, we selected 10 phylogenetically distinct retrons from our previous work that support higher levels of editing in *E. coli* to test in new bacterial species in this work (**Fig 1d**; Supplementary Table 2). We created a set of entry plasmids with each of the chosen 10 retron RTs and their corresponding retron ncRNAs. As the portability of the SSAP/SSB pairs has been analyzed in previous works (Filsinger et al., 2021), each plasmid additionally encodes an SSAP and compatible SSB for recombineering. When making single base edits, it is typical to suppress mismatch repair using *mutS* deletions or use of dominant negative MutL (Supplementary Fig 2). In this work, we tested sequential multi-base edits, which are not antagonized by mismatch repair, to remove that variable. In most species we used CspRecT as the SSAP, EcSSB as the SSB, and RSF1010 as the origin of replication. However, the SSAP, SSB, and origin of replication does differ for some species as discussed in the results to come.

To facilitate the cloning of editing donors, we used a single-pot Golden Gate workflow in which the set of entry plasmids for each species was mixed with a 70 bp dsDNA donor sequence encoding the edit to be made flanked by SapI golden gate ends (**Fig 1e**; Supplementary Fig 2). Individual colonies were screened to yield the 10 final plasmids for each species with the same donor but based on different retrons. Recombitron-containing plasmids were then introduced into cells using a range of mechanisms such as natural competence, electroporation or conjugation as appropriate for each species **(Fig 1e)**. For recombineering experiments, replicates of individual colonies were grown in liquid cultures, highly diluted in the presence of the inducer and grown again until saturation. Multiplexed amplicon sequencing (Illumina) was used to determine overall recombineering rates in the target locus **(Fig 1e)**.

### Retron-mediated recombineering in coliforms

We first extended retron recombineering technology to other coliforms, species closely related to *E. coli* **(Fig 2a)**. We selected two additional enterobacteria species in which oligonucleotide recombineering was previously tested and validated: *Citrobacter freundii* ATCC 8090 and *Klebsiella pneumoniae* ATCC 10031 (Wannier et al., 2020), and we compared these directly to *E. coli* bMS.346, an engineered host lacking *exo1* and *recJ* in which retrons produce the highest recombineering rates (Lopez et al., 2022). *C*. *freundii* has been used extensively in the biomanufacturing industry, but multiplexed engineering of metabolic pathways remains laborious (Jiang et al., 2012; Metsoviti et al., 2013). Recent studies have also revealed the role of *C. freundii* as a human pathogen that can cause gastroenteritis and urinary tract infections (Ramos-Vivas et al., 2020; Chen et al., 2023). *K. pneumoniae* was chosen because hypervirulent multidrug-resistant isolates cause pneumonia and other infections, representing one of the most serious threats to public health in the coming years (Tommasi et al., 2015; Tang et al., 2020).

**Figure 2.**
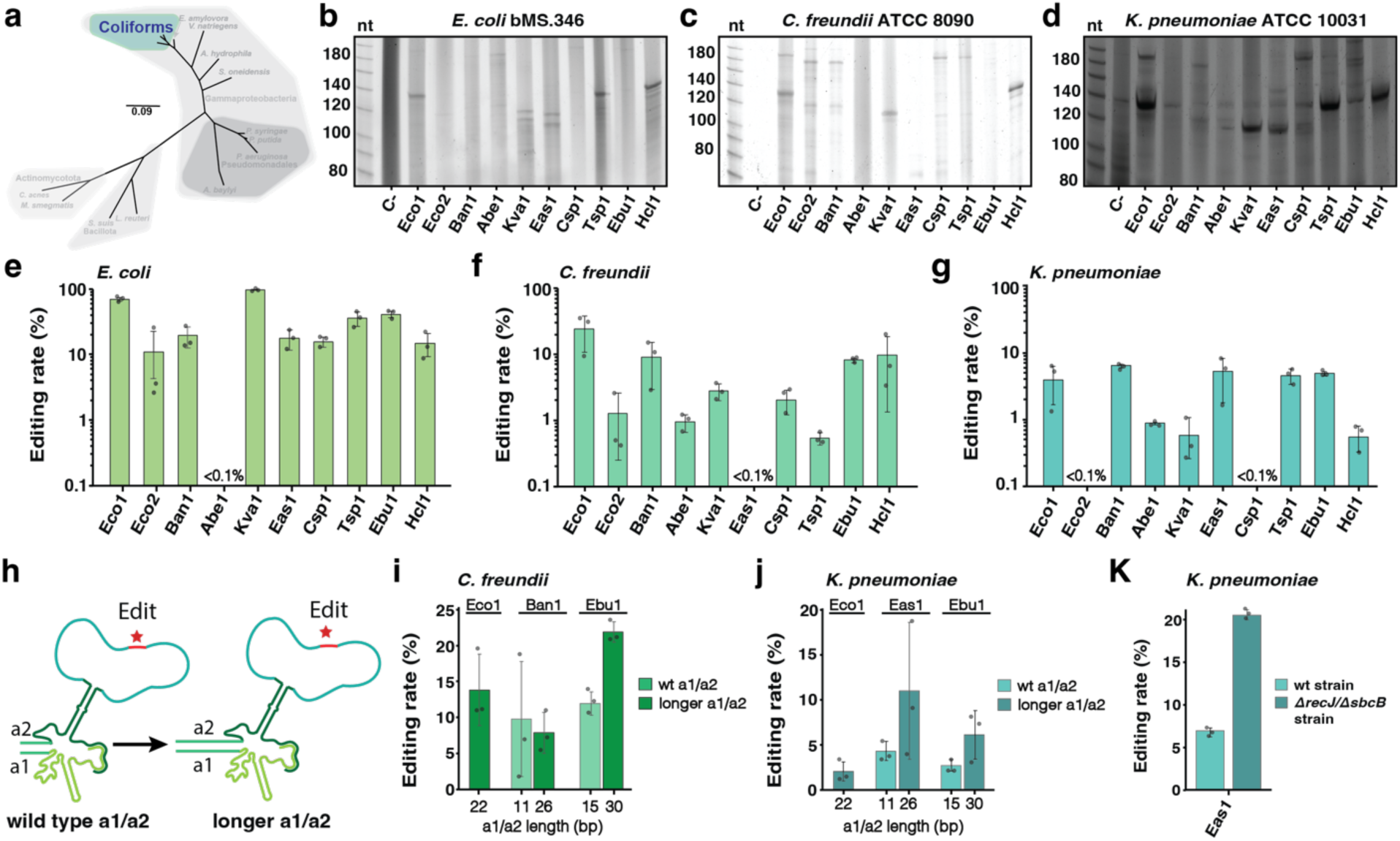
Retron-mediated recombineering in coliforms. **a**, Schematic of the phylogenetic tree from **Fig 1b**, highlighting the group of bacteria evaluated in this figure, the Coliforms. **b**, PAGE analysis of RT-DNA production in *E. coli* bMS.346. **c**, PAGE analysis of RT-DNA production in *C. freundii* ATCC 8090. **d**, PAGE analysis of RT-DNA production in *K. pneumoniae* ATCC 10031. In **b-d**, each RT-DNA was prepped from cells once and their length relative to a ssDNA ladder with markers in nt for nucleotides is indicated (uncropped gels are shown in Supplementary Fig 3). **e**, Quantification of precise genome editing to make a 5 bp deletion in the *lacZ* gene in *E. coli* using the 10 recombitrons set. **f**, Quantification of precise genome editing to make a 5 bp deletion in the *lacZ* gene in *C. freundii* using the 10 recombitrons set. **g**, Quantification of precise genome editing to make a 5 bp deletion in the *lacZ* gene in *K. pneumoniae* using the 10 recombitrons set. **h**, Schematic illustrating the lengthening of a1-a2 regions of retron ncRNA to boost editing rates. **i**, Quantification of precise genome editing in *C. freundii* using longer a1-a2 region in - Ban1 and-Ebu1 recombitrons in comparison with wild type versions. **j**, Quantification of precise genome editing in *K. freundii* using longer a1-a2 region in-Eas1 and-Ebu1 recombitrons in comparison with wild type versions. **k**, Quantification of precise genome editing with-Eas1 recombitron in wild type and *ΔrecJ/ΔsbcB* double knock-out *K. pneumoniae* strain. In **e-k**, data were quantified by sequencing after 24 h of editing using Illumina NextSeq, circles show each of the three biological replicates, and errors bars are mean ± standard deviation. Additional statistical details are presented in Supplementary Table 3.

To test retron-mediated recombineering in these coliform species, we designed recombitrons using the broad host range RSF1010 origin of replication and the high-performance SSAP, CspRecT, along with its compatible *E. coli* SSB (Wannier et al., 2020; Filsinger et al., 2021). Recombitron donors were designed to delete 5 bp in the *lacZ* gene for these species. For these initial experiments in coliforms, we checked for RT-DNA production from the recombitrons prior to quantifying editing. After inducing recombitrons for 5 hours in liquid cultures and isolating RT-DNA using a Qiagen miniprep followed by an ssDNA cleaning and concentration step, we visualized RT-DNA bands produced by each of the 10 recombitrons on TBE (Tris-Borate EDTA)-urea gels. We found substantial production for many retrons in all three species, evidenced by the ssDNA bands, with retrons-Eco1 and-Hcl1 yielding the highest production in all 3 species tested (**Fig 2b-d**; Supplementary Fig 3a-f). Interestingly, in this particular *K. pneumoniae* strain, an additional ssDNA band is apparent across all cultures including the negative control, which expresses no recombitron. We used DefenseFinder (Tesson et al., 2022) on the genome of this strain and found a putative type V retron reverse transcriptase gene which may explain the presence of this extra band.

We next tested recombineering in these species with induction of recombitrons for 16 to 24 hours. For context in our pre-engineered *E. coli* bMS.346, retron-Kva1 yielded editing of 97.3 ± 1.7 % of genomes (**Fig 2e**). This data confirms a previous result showing that the Kva1 recombitron outperforms an Eco1 recombitron in this bacterial strain for making single point mutations in the bacterial chromosome (Khan et al., 2024). In *C. freundii* ATCC 8090, the retron-Eco1 recombitron showed the highest editing rates (24 ± 13.5%), while recombitrons based on retrons-Ban1,-Ebu1 and-Hcl1 yielded ∼10% of genomes edited, and the rest resulted in lower editing rates (**Fig 2f**). In *Klebsiella pneumoniae* ATCC 10031, several recombitrons including those based on retrons-Ban1,-Eas1,-Tsp1 and-Ebu1 yielded editing rates around 5%, while the rest yielded lower rates (**Fig 2g**). To know whether promoter strength could increase editing rates, Pm, J23115 and lac promoters were tested with the Eco1 recombitron in the three coliforms, but showed no significant changes in recombineering efficiency between them (Supplementary Fig 3g-i). Taken together, these data extend retron recombineering beyond *E. coli* for the first time and reveal species-specific recombitron performance, highlighting the importance of screening for a set of retrons when a new species is tested.

It is important to note that the retron-Eco1 recombitron we used was an improved version with longer a1-a2 regions that increase RT-DNA production and editing rates in *E. coli* (**Fig 2h**; Lopez et al., 2022). To test whether this structural change could increase editing efficiencies in other retrons in other species, we tested lengthening the a1/a2 region in the best-performing recombitrons in *C. freundii* and *K. pneumoniae*, adding 15 bp to each of the a1/a2 ncRNA regions. In *C. freundii*, modification of the a1/a2 region in the retron-Ban1 recombitron did not affect editing, but extension of a1/a2 in the retron-Ebu1 recombitron doubled the editing efficiency, from 12.6 ± 1.4% to 21.9 ± 1.6% (**Fig 2i**). Similarly, in *K. pneumoniae* the longer a1/a2 regions nearly doubled retron-Eas1 and-Ebu1 recombitron editing rates, reaching numbers close to 10% in comparison with 5% when using the wild-type a1-a2 regions (**Fig 2j**). Finally, to test whether removal of *recJ* and *sbcB* would increase recombineering efficiencies in other species, as they do in *E. coli*, we knocked out these two genes in *K. pneumoniae.* Using the modified retron-Eas1 recombitron with longer a1-a2 region, these two mutations nearly tripled editing rates versus the modified recombitron in the wild-type strain, from 6.8 ± 0.2% to 22.2 ± 0.8% (**Fig. 2k**). Overall, these findings highlight the fact that strategies for editing optimization are portable across different retrons and different species.

### Retron-mediated recombineering in environmental Gammaproteobacteria

To further extend the use of recombitrons, we next tested them in environmental Gammaproteobacteria species where recombineering technology is rarely implemented, including *Erwinia amylovora* Eat8, *Vibrio natriegens* ATCC 14048, *Aeromonas hydrophila* ATCC 7966 and *Shewanella oneidensis* JG2150 **(Fig 3a)**. There are several reasons to choose these species. *E. amylovora* represents a threat to the food industry by causing fire blight, a disease affecting the production of apples, pears as well as other species in the *Rosaceae* family (Bonn et al., 2000; van der Zwet et al., 2012). *V. natriegens* has the fastest growth rate of any organism known, turning it into an increasingly popular chassis for molecular and synthetic biology purposes (Weinstock et al., 2016). *A. hydrophila*, a bacteria found in aquatic environments and well-known to be a fish pathogen, has recently drawn attention for causing an increasing number of diseases in humans such as wound infection, septicemia, urinary tract infections, and pneumonia (Janda and Abbott, 2010; Schwartz et al., 2024). Finally, *S. oneidensis’* versatile respiratory capacities, including utilization of organic and inorganic compounds, together with its specialized electron-transport pathways enable it to function as an electrochemically active bacteria (EAB) with potential applications in a variety of biotechnological processes including microbial fuel cells and generating CO_2_-free clean energy (Hau and Gralnick, 2007; Muñoz-Cupa et al., 2021; Yang et al., 2021).

**Figure 3.**
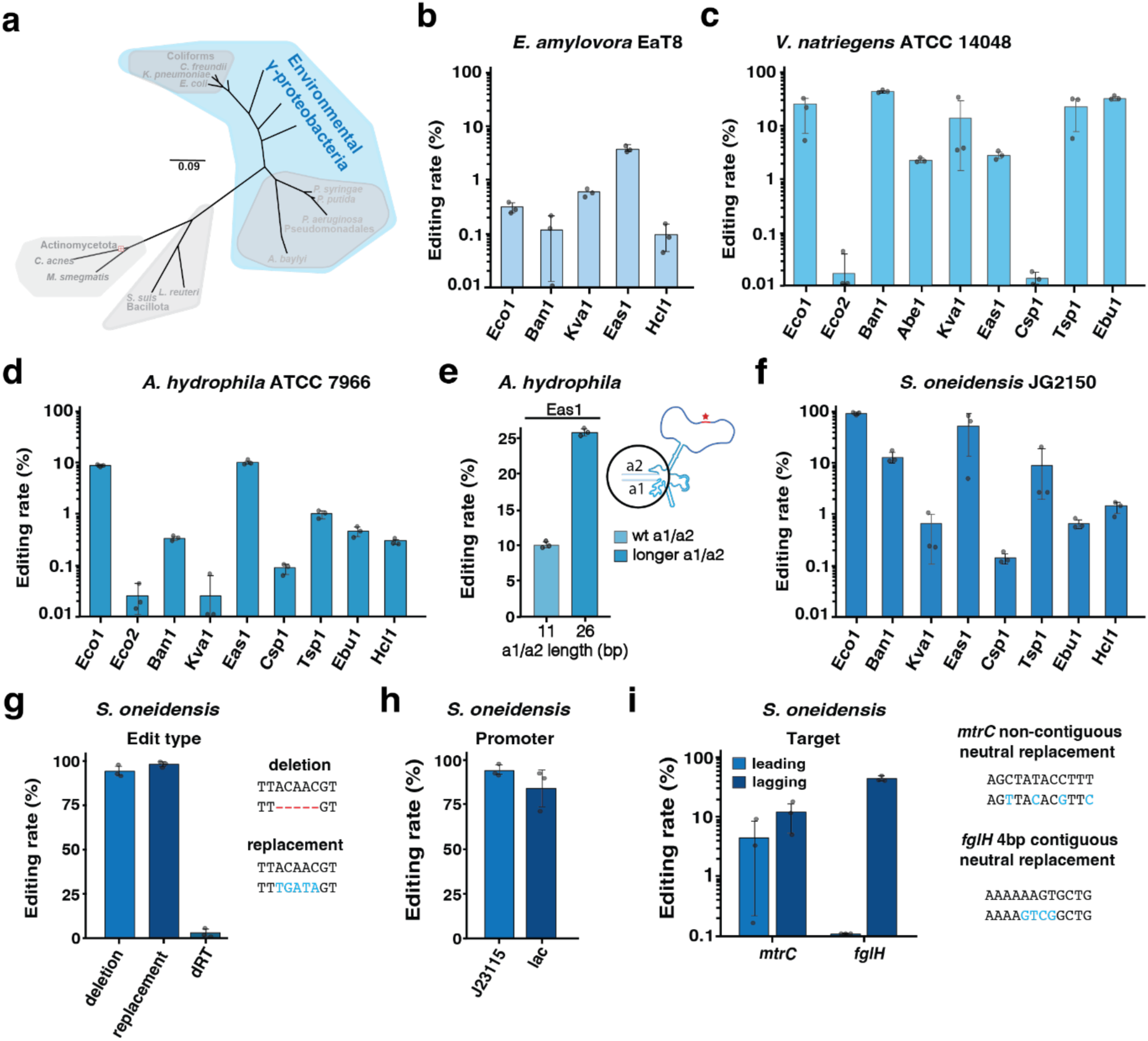
Retron-mediated recombineering in environmental *Gammaproteobacteria*. **a**, Schematic of the phylogenetic tree from **Fig 1b**, highlighting environmental Gammaproteobacteria. **b**, Quantification of precise genome editing to make a 5 bp deletion targeting a non-essential intergenic region in *E. amylovora* Eat8 using a set of 5 recombitrons. **c**, Quantification of precise genome editing to make a synonymous 4 bp replacement donor (GAGT to ATCG) targeting the *flgH* gene in *V. natriegens* ATCC 14048 using a set of 9 recombitrons. **d**, Quantification of precise genome editing to make a synonymous 4 bp replacement donor (TTCG to CAGT) targeting the *flgH* gene in *A. hydrophila* ATCC 7966 using the 10 recombitrons set. **e**, Quantification of precise genome editing in *A. hydrophila* using longer a1-a2 region with-Eas1 recombitron in comparison with wild type version. **f**, Quantification of precise genome editing to make a 5 bp deletion in the *lacZ* gene in *S. oneidensis* JG2150 using a set of 8 recombitrons. **g**, Quantification of precise genome editing to make different edits types in the *lacZ* gene in *S. oneidensis* JG2150 using-Eco1 recombitron. A dead version of Eco1 RT was used as a negative control. **h**, Quantification of precise genome editing to make a 5 bp deletion in the *lacZ* gene in *S. oneidensis* JG2150 using lac or J23115 promoter. **i**, Quantification of precise genome editing to make a non-contiguous neutral replacement in *mtrC* gene and a contiguous neutral replacement in *flgH* gene in *S. oneidensis* JG2150 with-Eco1 recombitron. In **b-i**, data were quantified by sequencing after 24 h of editing using Illumina NextSeq, circles show each of the three biological replicates, and errors bars are mean ± standard deviation. Additional statistical details are presented in Supplementary Table 3.

After electroporation or conjugation of recombitron plasmids into these species, recombineering experiments were carried as in coliforms. Beginning with *E. amylovora*, we designed a donor to make a 5 bp deletion targeting a non-essential intergenic region. In this species, only 5 out of 10 of the retrons tested produce any detectable edits and the retron-Eas1 based recombitron achieved the highest rate of 3.8 ± 0.5 % **(Fig 3b)**. In *V. natriegens* and *A. hydrophila*, a synonymous 4 bp replacement donor (GAGT to ATCG) targeting the *flgH* gene involved in flagella synthesis was used for editing experiments. In *V. natriegens* we tested 9 retrons with highly divergent results: whereas retron-Eco2 and-Csp1 recombitrons barely reached 0.01% editing, retrons-Eco1,-Kva1,-Tsp1 and-Ebu1 reached editing rates above 10%, and the retron-Ban1 recombitron achieved 44.7 ± 0.1 % of genomes edited **(Fig 3c)**. In *A. hydrophila*, the retron-Eas1 recombitron reached the highest editing rates of 9.9 ± 0.6 % **(Fig 3d)**. As longer a1/a2 regions boosted retron-Eas1 recombineering efficiencies in *K. pneumoniae* (**Fig 2j**), we also tried this improved architecture in *A. hydrophila* to show the portability of these modifications in phylogenetically distant species reaching 25.7 ± 0.4 % editing in the *flgH* gene **(Fig 3e).**

In *S. oneidensis,* we were able to use the same donor as we used for coliform experiments because the strain encodes an identical *lacZ* gene integrated in a single copy on its chromosome that was previously assayed to test oligonucleotide recombineering (Corts et al., 2019). Initial attempts at editing reached a technical failure point, where mutations in recombitron components or in the *xylS* repressor were detected by whole plasmid sequencing upon electroporation of the plasmids in this host. We modified our plasmids to test substituting the XylS/Pm inducible system with a constitutive J23115 promoter and used conjugation to introduce recombitron plasmids into *S. oneidensis*. We found that these constitutive versions did not accumulate plasmid mutations. Therefore, we ran editing experiments with these constitutively expressed versions of the ten recombitrons in *S. oneidensis*. The experiments were carried out in the same culture parameters and timing as for other Gammaproteobacteria, but the absence of uninduced recovery and outgrowth phases does mean that the recombitron was active in these cultures for a longer time. In *S. oneidensis*, the retron-Eco1 recombitron was the best-performer achieving 92.1 ± 4.8 % editing rate for a 5 bp deletion **(Fig 3f)**. As these numbers represent the highest recombitron efficiencies using a non-optimized strain, we decided to further investigate recombitron capabilities in this species. First, we measured retron-Eco1 editing rates for a 5 bp replacement rather than deletion, obtaining 97.9 ± 1.8 % efficiency **(Fig 3g)**. A dead RT version of Eco1 did not result in substantial editing, demonstrating that these high recombineering rates depend on the reverse transcribed donor **(Fig 3g)**. We also replaced the J23115 promoter with a lacI/pLac inducible system, which yielded 82.5 ± 10.1 % editing rates, showing that high-performance yielded by the retron-Eco1 recombitron in *S. oneidensis* is not solely due to the constitutive promoter, and could be achieved with an inducible promoter **(Fig 3h)**. Finally, to show the ability of Eco1 to edit different *loci* we tested recombitron efficiencies against several targets, including a non-contiguous neutral replacement in *mtrC* reaching editing rates up to 11.6 ± 6.9 %, and a contiguous neutral replacement in *flgH* gene with an efficiency of 44.2 ± 0.4 % **(Fig 3i)**.

### Retron-mediated recombineering in Pseudomonadales

Next, we tested recombitron technology in another relevant group among Gram-negative bacteria, the *Pseudomonales* order, which encompasses *Pseudomonadaceae* and *Moraxellaceae* families **(Fig 4a)**. We focused on three species from the *Pseudomonas* genus (*P. aeruginosa* PAO1, *P. putida* KT2440 and *P. syringae* B728a) in which oligo recombineering was previously tested with variable rates (Wannier et al., 2020; Swingle et al., 2010; Aparicio et al., 2020) and in *Acinetobacter baylyi* ADP1-ISx, a species with efficient homologous recombination where recombineering systems have not been tested to date (Elliot and Neidle., 2011). *P. aeruginosa* is a human opportunistic pathogen capable of causing a variety of infections in immunocompromised patients and it also shows a great ability to rapidly acquire resistance to multiple antibiotics constituting a challenge for modern medicine treatments (Angeletti et al., 2018; Sriramulu, 2019).

**Figure 4.**
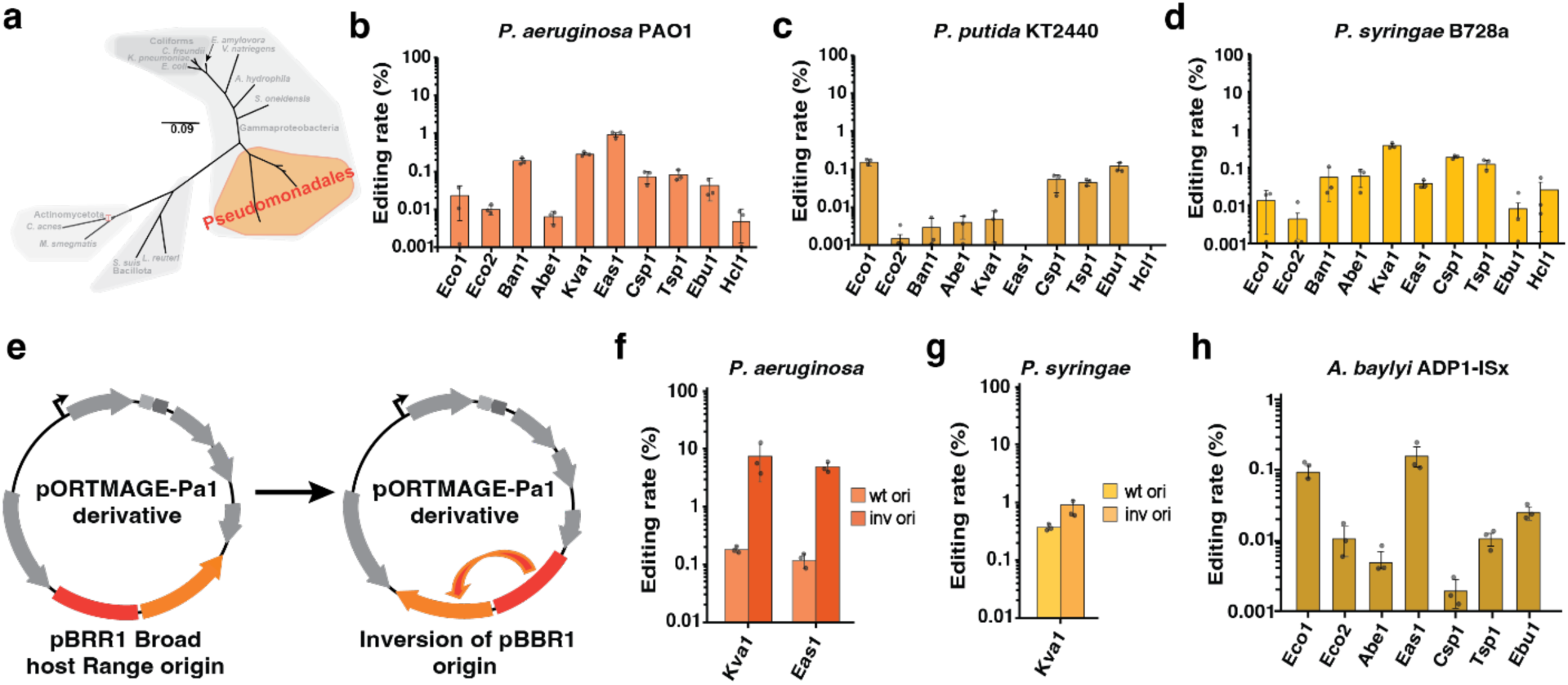
Retron-mediated recombineering in *Pseudomonadales*. **a**, Schematic of the phylogenetic tree from **Fig 1b**, highlighting the Pseudomonadales order. **b**, Quantification of precise genome editing to make a 4 bp deletion targeting the *phzM* gene in *P. aeruginosa* PAO1 using the 10 recombitrons set. **c**, Quantification of precise genome editing to make a 4 bp deletion targeting the *ykgJ* gene in *P. putida* KT2440 using the 10 recombitrons set. **d**, Quantification of precise genome editing to make a 5 bp deletion in a non-essential intergenic region in *P. syringae* B728a using the 10 recombitrons set. **e**, Schematic of the pORTMAGE-Pa1 derivative plasmids with wild type and inverted pBBR1 origin of replication. **f**, Quantification of precise genome editing using plasmids with wild type or inverted pBBR1 origin of replication in *P. aeruginosa* PAO1 using-Kva1 and-Eas1 recombitrons. **g**, Quantification of precise genome editing using plasmids with wild type or inverted pBBR1 origin of replication in *P. syringae* B728a using-Kva1 recombitron. **h**, Quantification of precise genome editing to make a 4 bp deletion targeting a glycosyltransferase gene in *A. baylyi* sFAB6437 using a set of 7 recombitrons. In **b-h**, data were quantified by sequencing after 24 h of editing using Illumina NextSeq. Circles show each of the three biological replicates, and errors bars are mean ± standard deviation. Additional statistical details are presented in Supplementary Table 3.

*P. putida* represents a versatile synthetic biology chassis due to its capacity for degrading toxic compounds and suitability for metabolic engineering (Loeschcke and Thies, 2015; Martínez-García and de Lorenzo, 2017). *P. syringae* is one of the most common plant pathogens, used as a model organism for host-pathogen interaction and one of the main threats for global crop production (Xin et al., 2018). Finally, *A. baylyi* exhibits a vast metabolic diversity, providing a strong basis for converting this host into a microbial cell factory in which extending the genome editing toolbox could lead to excel applications in synthetic biology (Santala and Santala, 2021).

We constructed all *Pseudomonas* recombitrons using the pORTMAGE-Pa1 backbone, a plasmid with pBBR1 broad host range origin of replication and PapRecT as SSAP (Wannier et al., 2020). Additionally, we added a compatible SSB (*P. aeruginosa* PaSSB) and removed mismatch repair dominant negative PaMutLE36K as we planned to make 4-5 bp edits in all the species. In *P. aeruginosa*, we designed a donor to make a 4 bp deletion in *phzM*, a gene involved in pyocyanin synthesis. After performing the retron-recombineering assay, the retron-Eas1 recombitron arose as the top performing, yielding a 0.92 ± 0.13 % editing rate **(Fig 4b)**. We also analyzed RT-DNA bands in *P. aeruginosa* on a TBE-Urea gel, which showed retron-Eas1 as well as-Tsp1 as top producers (Supplementary Fig 4a-b). In *P. putida*, the donor targets non-essential *ykgJ* gene to also make a 4 bp deletion. The results showed lower efficiencies in comparison with *P. aeruginosa* data, with retron-Eco1 recombitron as best editor, yielding 0.16 ± 0.02 % **(Fig 4c)**. Finally, in *P. syringae*, the donor was designed to make a 5 deletion in an intergenic region with no essential function. Here, the retron-Kva1 recombitron achieved 0.38 ± 0.02 %, the highest in this species **(Fig 4d)**.

Recombitrons tested in the three *Pseudomonas* species initially yielded much lower editing rates than other Gram-negative species tested. Although this is not entirely unexpected given lower editing rates were previously observed in these species using oligonucleotide recombineering (Wannier et al., 2020; Aparicio et al., 2020; Asin-García et al., 2023), we opted to investigate further to determine if any other retron-related factor was also limiting rates. It was recently demonstrated that a recombineering donor encoded in the retron *msd* on a plasmid can target itself if it is located in the lagging strand of the plasmid, competing with the target editing site and lowering the observed editing rate (Ni et al., 2024). To avoid this competition, the retron donor should be located in the leading strand of the plasmid. To test if plasmid competition for the donor may partially explain low editing rates in *Pseudomonas* species, we inverted the orientation of the pBRR1 origin of replication for the recombitron plasmids and repeated recombineering with these modified plasmids for all three species **(Fig 4e)**. Strikingly, editing rates were improved 8x in *P. aeruginosa*, with retron-Kva1 recombitron reaching 7.4 ± 4.7 % for the *phzM* target **(Fig 4f)**. In *P. syringae*, the efficiencies also improved 2x times using Kva1 recombitrons, respectively **(Fig 4g)**. Despite the success of this strategy for best editors, most retrons did not show any efficiency improvement (Supplementary Fig 4c-d), which may indicate a functional limitation of those retrons in *Pseudomonas* species.

For recombitron expression in *Acinetobacter baylyi*, we used pWBW162 (Biggs et al., 2020), a plasmid derived from high-copy pBAV1k broad host range vector previously shown to be propagated in *A. baylyi* (Bryksin et al., 2010). Recombineering has not previously been reported in *A. baylyi*, so we opted to use CspRecT/EcSSB to perform these experiments. The donor was designed to make a 5 bp deletion in a glycosyltransferase family protein gene. However, cloning the retrons in this backbone was challenging, and most colonies contained large deletions in the recombitron operon during the process of cloning in *E. coli*, indicating some incompatibility with retrons or CspRecT and this plasmid. After multiple attempts, we were able to obtain and transform a subset of four recombitrons into *A. baylyi* sFAB6437. However, editing rates were not able to reach efficiencies above 0.1% (Supplementary Fig 4e). Trying to solve the stability issue, we decided to clone the set of 10 retrons in the same RSF1010 origin of replication vector used in the coliforms and environmental gammaproteobacterial experiments. This alleviated issues with cloning and plasmid stability, and retron recombineering rates were slightly better than using the pBWB172 vector but still low, with recombitron-Eas1 as the top performer reaching 0.15 ± 0.08 % (**Fig 4h**; Supplementary Fig 4e).

### Retron-mediated recombineering in Bacillota and Actinomycetota

Next, we moved beyond the *Proteobacteria* phyla and tested recombitrons in two completely separate lineages within the bacterial domain, the phyla *Bacillota* and *Actinomycetota* **(Fig 5a)**. We chose two bacterial species from each phylum: For *Bacillota*, we chose *Limosilactobacillus reuteri* ATCC PTA 6475 (*L. reuteri* 6475) and *Streptococcus suis* P1/7 (*Bacillota*), and for *Actinomycetota*, we chose *Mycobacterium smegmatis* mc^2^ 155 and *Cutibacterium acnes* KPA17202. *L. reuteri* is a well-studied probiotic bacterium that produces antimicrobial molecules, which benefit the host immune system and strengthen the intestinal barrier by reducing translocation of pathogenic bacteria into other tissues (Mu et al., 2018; Yu et al., 2023). *S. suis* is an opportunistic zoonotic pathogen causing systematic disease in piglets with potential to cause disease in humans with close contact with pigs or exposed to contaminated pig meat (Wertheim et al., 2009; Fredrisken et al., 2024). *M. smegmatis* is a nonpathogenic and fast-growing species within the genus *Mycobacterium*, with thousands of conserved genes and similar physiology, representing an ideal model for mycobacterial research (Sparks et al., 2023). Finally, *C. acnes* is the most abundant skin commensal with some phylotypes linked to a healthy skin with others causing the development of acne vulgaris and other inflammatory diseases (Mayslich et al., 2021; Cros et al., 2023).

**Figure 5.**
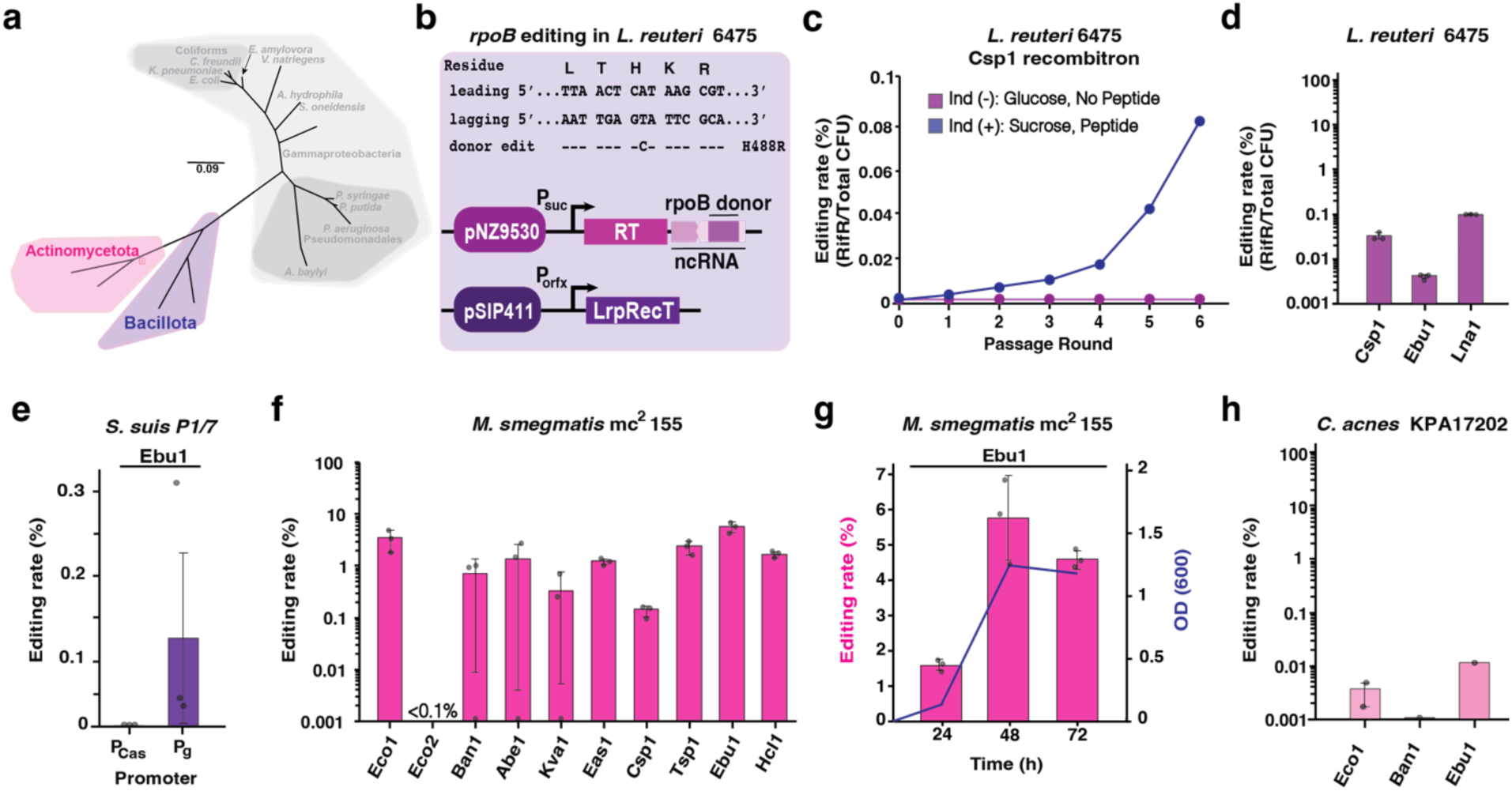
Retron-mediated recombineering in *Bacillota* and *Actinomycetota*. **a**, Schematic of the phylogenetic tree from **Fig 1b**, highlighting the *Bacillota* and *Actinomycetota* classes. **b**, Top: Schematic of the dsDNA sequence of the targeted *rpoB* region of *L. reuteri* 6475 is shown aligned with amino acids residues specified by each codon. On the left the leading and lagging strand are indicated and below the retron RT-DNA donor with the single-point mutation that results in a rifampicin-resistant phenotype. The amino acid change (H488R) is listed on the right. Bottom: Schematic of the tow plasmid assay used for retron-mediated recombineering in *L. reuteri*. One plasmid encoded the retron RT and engineered ncRNA to edit rpoB gene in the pNZ9530 backbone and under the P_suc_ promoter. The other plasmid encoded the LrpRecT gene under the P_orfx_ promoter in the pSIP411 backbone **c**, Quantification of precise genome editing to make a single-point mutation in *rpoB* (H488R) gene in *L. reuteri* in the absence or the presence of sucrose and inducing peptide with-Csp1 recombitron. The experiment was performed for 6 passage rounds. **d**, Quantification of precise genome editing to make a single-point mutation in *rpoB* (H488R) gene in *L. reuteri* using a set of 3 recombitrons following single passage. In **c-d**, editing rates were calculated as % of rifampicin resistant colonies respect to total number of colonies in no-antibiotic plates **e**, Quantification of precise genome editing to make a 5 bp deletion in *purA* gene in *S. suis* P1/7 using-Ebu1 recombitron expressed under P_Cas_ or P_g_ promoters. **f**, Quantification of precise genome editing to make a 5 bp deletion in MSMEG_5894 gene in *M. smegmatis* MC^2^ 155 using the 10 recombitrons set. **g**, Quantification of precise genome in *M. smegmatis* after 24, 48 and 72 h of editing and correlation with cell growth state (OD_600_) using-Ebu1 recombitron. **h**, Quantification of precise genome editing to make a 4 bp deletion in beta-galactosidase gene in *C. acnes* KPA17202 using the 10 recombitrons set. In **d-h**, data were quantified by sequencing after 24, 48 or 72 hours of editing using Illumina NextSeq, circles show each of the three biological replicates (except in C. acnes in which only one or two biological replicates were performed), and errors bars are mean ± standard deviation.

Despite the general lack of molecular tools that work in many Gram-positive bacteria, oligonucleotide recombineering technology has been characterized in lactic acid bacteria with relatively high efficiency in *L. reuteri* and *Lactococcus lactis* (van Pijkeren and Britton, 2012). To test recombitrons in *L. reuteri* 6475, we modified the pJP042, pSIP411 vector encoding the *recT1* gene, recently renamed as LrpRecT (Wannier et al., 2021), and cloned codon optimized retron components under the inducible P_orfx_ promoter (Risøen et al., 2000). We designed a 80 bp donor to target the RNA polymerase gene *rpoB* and place a C to A mismatch mutation, producing an amino acid change (H488R) and conferring rifampicin resistance in *L. reuteri* **(Fig 5b)**. Unfortunately, we found persistent, distinct mutations in either the RT or LrpRecT genes appearing after transformation of these plasmids when cloning parts in *E. coli*. Only the retron-Csp1 recombitron was able to be constructed with no mutations in *E. coli*. However, it became unstable when transformed into *L. reuteri* 6475. To reduce toxicity due to the recombineering components, we cloned the retron-Csp1 recombitron into low copy vector pNZ9530 (Pavan et al., 2000) under P_suc_ promoter, a sucrose inducible promoter. After obtaining a transformant with no mutations, we assayed recombineering efficiency during 5 passages in *L. reuteri* 6475, but found no increase in the abundance of the intended edit over time (Supplementary Fig 5a). Furthermore, at the end of the passages, we found dominant mutations in the RT and LrpRecT, indicating that the plasmid in *L. reuteri* 6475 cannot stably maintain the full recombitron operon.

To overcome this issue, we next constructed a two-plasmid system: the retron-Csp1 recombitron was cloned in pNZ9530 under P_suc_ promoter and pVPL3017, pJP042 erythromycin resistant marker swapped with chloramphenicol resistant marker, was used for LrpRecT expression (Oh and van Pijkeren, 2014) **(Fig 5b)**. Using the two-plasmid system, the abundance of the intended edit went up with every passage in the induced condition, but not in the uninduced condition, reaching ∼ 0.1% after 6 passages **(Fig 5c)**. Rifampicin-resistant colonies were used to confirm targeted mutations in the *rpoB* gene using mismatch amplification mutation assay (MAMA) PCR (Cha et al., 1992; Supplementary Fig 5b). The low editing rates observed after 6 passages of the experiment could be due to a lack of activity of the retron-Csp1 recombitron in this species or by negative fitness of the mutation selecting against edited cells. Indeed, growth assays showed that *rpoB* mutants were impaired relative wild-type cells (Supplementary Fig 5c). Given the positive results with a two-plasmid system in *L. reuteri*, we cloned and tested two additional retrons in the pNZ9530 vector, retron-Eco1 and-Ebu1. Additionally, to test whether a more adapted retron could be more active in this host, we also cloned a retron known as Lna1, originally from *Lactobacillus nantensis*, which had shown some editing efficiency in a previous work in *E. coli* (Khan et al., 2024). Retron-Ebu1 and Lna1 were successfully cloned in *L. reuteri* 6475 while retron-Eco1 was not maintained in pNZ9530. Three recombitrons (Csp1, Ebu1, and Lna1) were stably maintained in pNZ9530 in *L. reuteri* 6475 with no mutations after single passage following induction. Interestingly, retron-Lna1 showed about 0.1% recombineering efficiency after single passage, while retron-Csp1 and-Ebu1 yielded less than 0.01% recombineering efficiency **(Fig 5d)**. However, retron-Lna1 did not improve recombineering efficiency following multiple passages (data not shown) possibly due to the instability of plasmid or potential mutations in retron-Lna1 in *L. reuteri* 6475.

Although there are no prior reports of recombineering in *S. suis*, a recent study used a dsDNA repair template plus Cas9 counterselection to precisely engineer the genome of this organism (Gussak et al., 2023). We adapted the pSStarget vector used in that work to construct the plasmids expressing the recombitron operon. As no *recT* gene has been experimentally reported for this host we used CspRecT and EcSSB for the recombineering assay. We constructed 8 recombitrons to encode a 70 bp donor to make a 5 bp deletion in *purA*, a gene involved in purine nucleotide synthesis. The plasmids were placed in *S. suis* P1/7 using a natural competence protocol (Zaccaria et al., 2014). However, no editing was detected with any of the constructs tested. To tackle this challenge in *S. suis* P1/7 we focused in a single recombitron, retron-Ebu1. To enhance Ebu1 expression, we replaced P_Cas_, the Cas9 native promoter with P_g_, a strong constitutive promoter that drives the expression of the housekeeping gene *gadpH* and has been shown to have the highest activity in this species (Zhang et al., 2022). We repeated the experiment with the new construct obtaining editing rates of 0.13% ± 0.12 % using Ebu1 recombitron **(Fig 5e)**. This result highlights the occasional need to optimize multiple elements involved in recombineering such as the promoter, vector, and SSAP/SSB pairs in addition to the retron to obtain detectable editing in the new species tested.

In contrast to *Streptococcus*, oligonucleotide recombineering has been previously used in *Mycobacterium* (van Kessel and Hatfull, 2007; van Kessel and Hatfull, 2008). To edit *M. smegmatis* mc^2^ 155, we used the pJV62 vector encoding mycobacteriophage Che9c gp61, recently renamed as MspRecT (Wannier et al., 2021), to construct the recombitron plasmids. The 10 retron operons were codon optimized and cloned upstream of the MspRecT with an engineered ncRNA containing a donor to make a 5 bp deletion that knocks-out MSMEG_5894, a gene coding for Mam4a protein which is essential for mycobacteriophage Brilliant infection (unpublished data). Moreover, MSMEG_5894 knockout bacteria cannot grow on minimal medium (MM) plate with cholesterol as the carbon source, giving multiple methods to quantify successfully edited colonies. After transformation of the recombitron plasmids into *M. smegmatis,* single colonies were grown in liquid culture until saturation followed by a 1:1000 dilution, induction with acetamide and succinate and sample collection every 24 hours for 3 days. After 24 hours, we were able to detect edited colonies in MM with cholesterol and colonies resistant to phage Brilliant infection (Supplementary Fig 5d-e). In general, multiplexed sequencing results show that the highest editing rates were achieved after 48 hours of induction and the top editor was the retron-Ebu1 recombitron reaching 5.7 ± 1.3 % efficiency **(Fig 5f)**. Nevertheless, Ebu1 performance did not increase after 72 hours, directly correlating with a plateau in the OD_600_ of the culture, showing again that recombitron editing is dependent on replication **(Fig 5g)**.

Finally, we evaluated retron-mediated recombineering in *C. acnes* KPA171202, a bacterium in which only homologous recombination has been reported to date (Sörensen et al., 2010; Knödlseder et al., 2024). Recombitrons encoding the RT, a ncRNA with a donor to make a 4 bp deletion in a beta-galactosidase gene, and the CspRecT/EcSSB pair were cloned under the p1340 promoter in vector pBRESP36A (Jore et al., 2001). Since *C. acnes* is a slow-growing, anaerobic bacterium, the experiment experimental pipeline had to be adjusted. Selected colonies were replated and grown anaerobically for another 48h before being inoculated to a starting OD of 0.1, re-inoculating every 24h for a minimum of three passages. No editing was detected with any of the constructs after amplicon sequencing. To make recombitrons work in this host, we looked for the presence of *recT* genes in phage genomes infecting related bacteria. We found *recT* genes in *P. freudenreichii* phages Anatole and Doucette (Modlin et al., 2016a; Modlin et al., 2016b), that we named as AnaRecT and DouRecT, respectively (Supplementary Fig 5f). We constructed new plasmids for Eco1, Ban1 and Ebu1 recombitrons encoding AnaRecT or DouRecT coupled with *C. acnes* SSB (CaSSB). After the recombineering experiment was performed, the retron-Ebu1 recombitron combined with AnaRecT show a low recombineering efficiency around 0.01% for a single replicate and **(Fig 5h)**. Thus, further efforts will be required to make recombitron technology practical in *C. acnes*.

## DISCUSSION

In this work we survey the portability and versatility of retron-mediated recombineering across three different bacterial phyla and a total of 15 different species, demonstrating the capacity of this novel technology to precisely edit prokaryotic genomes beyond the model organism *E. coli*. The involvement of nine research labs to perform this comprehensive survey shows that coordinated efforts can accelerate the development and spread of new genome editing tools, paving the road to other labs to rapidly adopt a new technology worldwide, a process which normally takes many years. Moreover, the distributed nature of the experiments across multiple sites further demonstrates the portability of the approach not just across species, but across different experimenters working under the slightly different experimental settings that occur across different labs. Taking advantage of the strong background of every research lab in the species tested, we show that recombitrons work in all of the hosts, with editing rates ranging from 0.01% to 98% **(Table 1)**. Furthermore, we also demonstrate that previous improvements made to recombitron parts, such as extending the a1/a2 regions (Lopez et al., 2022), are also portable between species leading to enhanced editing rates in some of the assessed organisms. Strikingly, recombineering efficiencies above 40% were reached in *V. natriegens* and *S. oneidensis* with no additional optimization, demonstrating that there are species where the technology is immediately ready for deployment to answer biological questions.

**Table 1.** Best retron editor in each host used in this study.

| Species | SSAP | Best Retron | Retron Editing rate | Highest reported oligo AFR | oligo editing ref. |
| --- | --- | --- | --- | --- | --- |
| *Escherichia coli* | CspRecT | Kva1 | 97.4% | 51% | Wannier et al., (2020) |
| *Citrobacter freundii* | CspRecT | Ebu1 | 21.9% | 36% | Wannier et al., (2020) |
| *Klebsiella pneumoniae* | CspRecT | Eas1 | 22.2% | 11% | Wannier et al., (2020) |
| *Erwinia amylovora* | CspRecT | Eas1 | 3.8% | ND |  |
| *Vibrio natriegens* | CspRecT | Ban1 | 44.7% | ND | Lee et al., (2019) |
| *Aeromonas hydrophila* | CspRecT | Eas1 | 25.7% | ND |  |
| *Shewanella oneidensis* | CspRecT | Eco1 | 98.0% | 5% | Corts et al., (2019) |
| *Pseudomonas aeruginosa* | PapRecT | Kva1 | 7.4% | 15% | Wannier et al., (2020) |
| *Pseudomonas putida* | PapRecT | Eco1 | 0.2% | 5.8 | Aparicio et al., (2020) |
| *Pseudomonas syringae* | PapRecT | Kva1 | 0.7% | 0.024% | Swingle et al., (2010) |
| *Acinetobacter baylyi* | CspRecT | Ban1 | 0.15% | ND |  |
| *Lactobacillus reuteri* | LrpRecT (RecT1) | Lna1 | 0.10% | 11% | van Pijkeren et al., (2012) |
| *Streptococcus suis* | CspRecT | Ebu1 | 0.13% | ND |  |
| *Mycobacterium smegmatis* | MspRecT (gp61) | Ebu1 | 5.7% | 0.3% | Filsinger et al., (2025) |
| *Cutibacterium acnes* | AnaRecT | ND | 0.015% | ND |  |

ARF, allelic recombination frequency; ND, Not Determined

The results obtained in the present survey show that retron-mediated editing outperforms oligonucleotide recombineering in five of the nine species where oligonucleotide recombineering has previously been evaluated **(Table 1)**. We also show that recombitrons work in six new species in which no recombineering technologies have been successfully demonstrated **(Table 1)**. In *C. freundii* and *P. aeruginosa* recombineering rates were very similar to those reached using oligonucleotides **(Table 1)**. However, oligo recombineering is clearly better in P. putida and L. reuteri **(Table 1)**. We expect that additional fine-tuning of recombitron parts in these hosts would enhance editing efficiency beyond oligonucleotides.

We observed low functional editing in one species, *C. acnes*, even with the use of phylogenetically related SSAPs (AnaRecT and DouRecT), signaling that broader efforts are needed in this species to enable retron recombineering. In this regard, a recent study using a library screening approach to test divergent SSAPs in 6 bacterial species has shown that the most effective SSAPs frequently originated from phyla distinct from their bacterial hosts (Filsinger et al., 2025). Future efforts to test a set of RecTs and retrons in parallel could facilitate the implementation of recombineering tehcnologies in previously uncharacterized bacterial species.

Interestingly, the particular retron that led to the highest editing differed substantially across species, highlighting the importance of testing several editors when a novel host is assayed **(Figure 6)**. For instance,-Kva1 recombitron is the top performer in *E. coli*, but is not even close to the top editor in the phylogenetically related *C. Freundii* and *K. pneumoniae* **(Fig 2d-f**, **Fig 6)**. In general, retron-Kva1,-Ban1 and retrons belonging to clade 9 are the best performers in most bacteria **(Fig 6)**.

**Figure 6.**
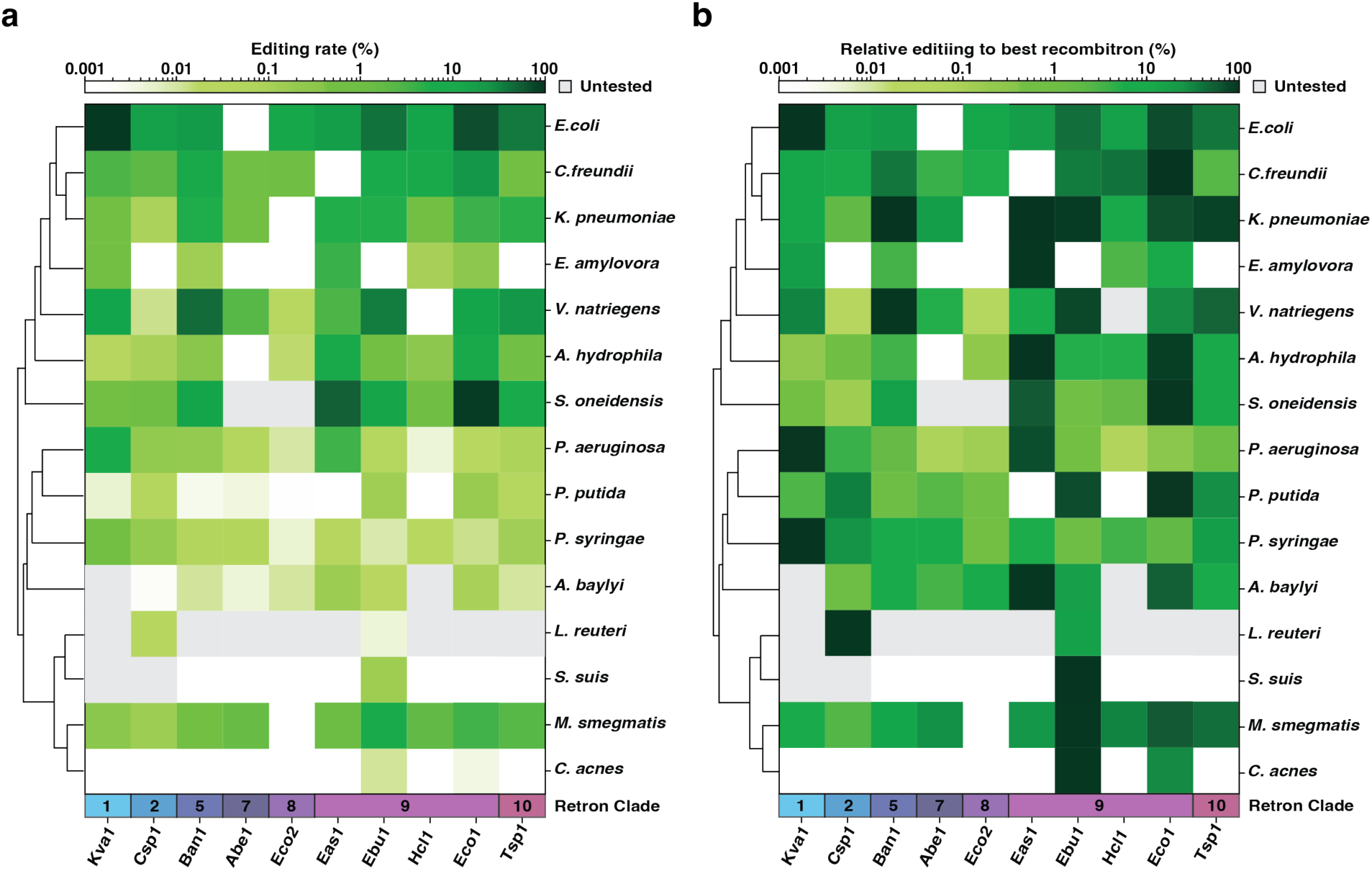
Summary of retron-mediated recombineering across the Bacteria domain. **a**, Heatmap depicting the absolute value of retron-mediated recombineering efficiency in the 15 bacterial species tested. **b**, Heatmap depicting the value of retron-mediated recombineering efficiency normalized to the best recombitron in each one of the bacterial species aasayed. In **a-b**, logarithmic scale was used to show the compiled editing data from **fig 2-5** in a scale of greens (light green, low editing; dark green, high editing), untested recombitrons were shown in grey color and no editing recombitrons were shown in white. On the left a tree indicates the phylogenetic relationships of the bacterial species tested in the work. The retron clade (from Mestre et al., 2020) is shown above of the retron names.

In *E. coli*, the first description of retron recombineering reported an editing rate of <0.1% (Farzadfard et al., 2014), but since that time both the cell and the retron have been engineered to achieve enhanced editing rates of >95% as reported here and in other contemporaneous studies (Liu et al., 2023; Ni et al., 2024). These modifications include lengthening the a1-a2 regions in the ncRNA (Lopez et al., 2022), inverting the origin of replication of the recombitron vector (Ni et al., 2024), and pre-engineering the host to knock-out *recJ* and *sbcB* genes (Schubert et al., 2021). To avoid host engineering which remain laborious and time-consuming, a CRISPRi strategy to transiently knockdown *recJ* and *sbcB* genes has also been used to boost retron-mediated recombineering in *E. coli* (Farzadfard et al., 2021). We expect that researchers working on the new species in this work that have detectible, but low, editing rates will continue to develop the technology to achieve such enhanced editing in their own chosen species.

It may be useful to contextualize retron recombineering in relation to other precise bacterial editing technologies, highlighting which technology is best suited for different types of edits. For instance, homologous recombination allows scarless and site-specific gene-size replacements, deletions or insertions of DNA sequences (Hamilton et al., 1989). In combination with CRISPR-Cas9 counterselection, homologous recombination has enabled large modifications across a wide range of bacterial species (Jiang et al., 2013; Mougiakos et al., 2017; Walker et al., 2020; Adalsteinsson et al., 2021; Stukenberg et al., 2022). However, the counterselection that is necessary to make homologous recombination efficient is often not practical to implement with smaller mutations that often have no potential gRNAs in the region or only a very limited number, making counterselection escape the most likely outcome. For single base substitutions, base editors—a fusion of Cas9 nickase with an adenine or a cytidine deaminase—represent a powerful platform with high efficiency in a variety of bacterial genomes (Tong et al., 2019; Xia et al., 2020; Volke et al., 2020). However, the deaminase editors are limited to single base pair substitutions and cannot make insertions and deletions, and additionally require a fortunate placement of both a PAM motif and deamination target for precise editing, which is often not present. Retron recombineering represents a versatile alternative to generate scarless modifications of moderate size, from single-nucleotides to small gene-sized insertions, deletions, or replacements (Ellis et al., 2001; Wang et al., 2009).

Retrons have been used not only for prokaryotic genome editing but also for other biotechnological applications, including molecular recording, phage editing, transcription factor modulation, and more. This work should also help these applications to be ported to a wider set of species. For instance, retron-based molecular recorders (Farzadfard et al., 214; Battharai-Kline et al., 2022) in envorimental or pathogenic bacteria could help researchers understand complex biological processes by creating noninvasive sentinel cells in specific niches. Phage editing beyond coliphages could accelerate the study and production of viral particles for phage therapy (Fishman et al., 2023). Reprogrammed retron-derived DNA for allosteric transcription factor dynamic regulation (Lee and Kim, 2023) could lead to advances in synthetic biology for modular, rapid, and post-translational control of protein localization in different microbes. Therefore, this work together with the previous effort to develop retron-based technologies should form a basis for the deployment of new technologies not only for bacterial genome editing but also for broader applications in a wide variety of bacterial species.

## METHODS

Biological replicates were taken from distinct samples, not the same sample measured repeatedly.

### Phylogenetic tree construction

The organisms used in this work are listed in Supplementary Table 1. The genome sequence of these organisms was obtained from the NCBI database, except for *Klebsiella pneumoniae* ATCC 10031 where the genome was obtained from American Type Culture Collection (ATCC) and *Erwinia amylovora* Eat8 where the genome was determined by in-house sequencing in the Koskella lab. For phylogenetic comparison, the 16S sequences were extracted from the whole genomes and aligned using MAFFT (Katoh and Standley, 2013) and the tree was constructed with FastTree 2 program (Price et al., 2010) with the WAG evolutionary model, and the discrete gamma model with 20 rate categories.

### Bacterial strains for cloning purposes

The *E. coli* strain used for cloning most plasmids used in this work was DH5α (New England Biolabs). To avoid mutations in any of the recombitron parts, *E. coli* strain ABLE C (Agilent) was used to reduce the copy number of p1340 vector (shuttle vector for *C. acnes*) by 4x fold. *E. coli* WM3064 strain (*thrB1004 pro thi rpsL hsdS lacZ*ΔM15 RP4–1360 Δ(*araBAD*)*567* Δ*dapA1341*::[*erm pir*(wild type)]) (Saltikov and Newman, 2003). The three strains were grown in LB medium (10 g l−1 tryptone, 5 g l−1 yeast extract and 5 g l−1 NaCl). Antibiotics were added as required (kanamycin, gentamycin, erythromycin and chloramphenicol).

### Recombitron-tested bacterial strains

All bacterial strains in which retron-mediated recombineering was evaluated are listed in Supplementary Table 1. Growth conditions of the different bacterial species used in this work are described in the following sections.

### Plasmid construction

All the plasmids used in this study are listed in Supplementary Table 4, and a subset is available from Addgene. Moreover, all the gBlocks (IDT) encoding the desired mutations for the retron-mediated recombineering experiments are listed in Supplementary Table 5. All PCR amplifications step were carried out with Q5-High Fidelity Master Mix (NEB). The plasmids used to test the optimized recombitron architectures were based on previously published pAGD229 plasmid (Gonzalez-Delgado et al., 2024). CspRecT was amplified from pORTMAGE-Ec1 vector (Wannier et al., 2020; Addgene #138474) and cloned downstream of the Eco1 retron operon in pAGD230 to create pAGD383. A KLD reaction (NEB) was used to replace T7 promoter with lac and J23115 promoters to generate pAGD439 and pAGD440 plasmids, respectively. The minimal recombitron operon (RT, retron ncRNA and CspRecT) encoded in these plasmids was cloned into a pET21 (+) vector (Novagen) to create pAGD383 with pBR322 origin of replication. To facilitate the description of the steps followed for the construction of plasmids to test the portability of retron-mediated recombineering, we grouped bacterial species based on the parental plasmid that was used to clone the set of 10 recombitrons.

#### Coliforms and environmental Gammaproteobacteria

In the seven bacterial species corresponding to these groups all the plasmids constructed were based in pORTMAGE-Ec1 vector. In a previous work, this vector was used to construct pAGD201 plasmid, which encodes Eco1 recombitron with dominant negative EcMutLE32K (Gonzalez-Delgado et al., 2024). To test multi-base edits, 4-5 bp deletions or replacements, the MutLE32K gene was replaced with EcSSB. EcSSB was amplified from the *E. coli* bMS.346 genome and cloned into pAGD201 plasmid generating pAGD322 (Supplementary Table 4). The set of 10 retrons used in this study (Supplementary Table 2) were individually amplified from plasmids constructed in a previous work (Khan et al., 2024), and resulting amplicons were run in 2% E-Gel EX (Invitrogen). The 10 amplicon bands were purified together and cloned into pAGD322 backbone with the RSF1010 origin of replication using a NEBuilder HiFi DNA assembly workflow (NEB). After transformation in the cloning strain, multiple colonies were screened using Sanger Sequencing to find the 10 recombitrons plasmids. The next step was using primers containing the SapI stuffer to amplify the newly constructed recombitrons. Again, the 10 amplicons were purified together followed by a KLD reaction, whose product was used to transform into the cloning strain. Multiple colonies were screened searching for stuffer-containing recombitrons. For experiments in environmental Gammaproteobacteria species XylS/Pm system was replaced with the lacI/P_lac_ system or J23115 constitutive promoter at this step using a Gibson Assembly reaction. A one-pot Golden Gate reaction (Engler et al., 2008) was used to substitute the SapI stuffer with the desired donor for each one of these hosts **(Fig 1e)**. An example of the method followed to clone a donor in the SapI stuffer region of Eco1 ncRNA is detailed in Supplementary Fig 2. The Golden Gate reaction was prepared in a volume of 20 µL as follows: 125 ng of SapI flanked donor (order as gBlock from IDT), 50 ng of a mix of the 10 recombitron plasmids from the previous step, 1.5 µL of SapI (NEB), 0.5 µL of T4 DNA ligase, 2 µL of T4 DNA ligase buffer and completed with 10 µL of H_2_O. The reaction consists of 30-60 cycles of 5 minutes at 37°C (for SapI) and 5 minutes at 16°C (for T4 DNA) ligase with a final step at 60°C for enzyme inactivation. 2-5 µL of the reaction were used to transform into cloning strain. Multiple colonies were screened using Sanger sequencing to find donor-containing recombitrons that were ready for performing the retron-mediated recombineering experiments.

#### Pseudomonas species

plasmids used in the three *Pseudomonas* species tested in this work were based in pORTMAGE-Pa1 vector (Wannier et al., 2020; Addgene #138475). Just like the preceding paragraph, PaSSB was amplified from *P. aeruginosa* PAO1 chromosome and cloned in pORTMAGE-Pa1, replacing PaMutLE36K to generate pAGD477 (Supplementary Table 4). The RT and SapI stuffer-containing ncRNA generated for coliforms and environmental Gammaproteobacteria were amplified and cloned using a Gibson Assembly approach upstream of the PapRecT gene to generated the set of recombitrons for *Pseudomonas* species. After screening for the different recombitrons using Sanger Sequencing, the Golden Gate reaction described in the preceding section was used to clone the desired donors for the experiments in the three hosts.

#### Acinetobacter baylyi

the full recombitron operons containing the SapI stuffer for coliforms and environmental Gammaproteobacteria were cloned into pBW162 vector (Addgene #140632) downstream of the trc promoter using a Gibson assembly reaction. The full set of recombitrons was verified using Sanger or whole plasmid sequencing. The above-described Golden Gate reaction was performed to clone the donor to edit *A. baylyi* into the retrons ncRNAs.

#### Lactobacillus reuteri

For inducible LrpRecT expression, *L. reuteri* 6475 was transformed with pVPL3017 (Cm^R^) (Oh and van Pijkeren, 2014). The recombitrons were codon-optimized for *L. reuteri* and synthesized (gBLOCK, IDT). Synthetic recombitrons were ligated with *L. reuteri* sucrose-inducible promoter (P_suc_) and inserted in pNZ9530 (Em^R^) by Ligase Cycling Reaction (LCR) (de Kok et al., 2014). The *L. reuteri* 6475 harboring pVPL3017 was directly transformed with LCR ligates (pNZ9530 containing recombitron cassettes) to establish two plasmids retron system.

#### Streptococcus suis

pSStarget vector (Gussak et al., 2023) was used as a backbone to clone the set of 10 recombitrons for *S. suis* experiments. The coliform-based recombitrons were cloned using Gibson Assembly downstream of the pCas promoter replacing the Cas9 operon, the ccdAB toxin/antitoxin system and the tetracycline resistance gene encoded in the parental vector. After sequencing verification, *S. suis* donor was cloned in the SapI stuffer using the one-pot Golden Gate reaction previously described. Pg promoter was order as gBlock (IDT) and cloned in the place of the Cas9 regulatory region in-Ebu1 recombitron (pLBL21) using Gibson Assembly to generate pLBL23 plasmid (Supplementary Table 4).

#### Mycobacterium smegmatis

pJV62 vector (Addgene #26910), which already encoded MspRecT, was used to construct the recombitron plasmids for this host. Codon optimized gBlocks (Twist Bioscience) with the set of 10 retron RTs with the ncRNA containing the donor to edit MSMEG_5894 were cloned into the pJV62 vector downstream of the acetamidase inducible promoter using a Gibson Assembly reaction.

#### Cutibacterium acnes

pBRESP36A vector (Jore et al., 2001) was used as backbone to clone codon optimized recombitrons. Codon optimized CspRecT and EcSSB were cloned downstream of the p1340 promoter using Gibson Assembly to create pAGD684 (Supplementary Table 4). Then, the 10 codon optimized RTs with ncRNA containing a donor targeting the C. acnes beta-galactosidase gene were cloned in pAGD684 upstream of the CspRecT/EcSSB pair. Codon optimized AnaRecT and DouRecT combined with CaSSB were cloned into pBRESP36A vector using Gibson Assembly, generating pAGD735 and pAGD738 plasmids, respectively.-Eco1,-Ban1 and-Ebu1 recombitrons were the recombitrons cloned upstream of AnaRecT and DouRecT.

### Electrotransformation of plasmids into bacterial species

#### Coliforms and Pseudomonas putida

The same electrotransformation protocol was used for *E. coli* bMS.346, *C. freundii* ATCC 8090, *K. pneumoniae* ATCC 10031 and *P. putida* KT2440. 100 ml of these four species were grown in LB medium at 37°C for coliforms and 30°C for *P. putida* until saturation. Cells were put on ice for approximately 10 min, washed three times with cold water, and resuspended in 1/100^th^ culture volume of cold water. 0.1-1 µg of recombitrons plasmids were mixed with 100 µL cell aliquots for 15 minutes on ice. This mixture was transferred to a 0.2-cm gap cuvette and electroporated immediately using a Gene Pulser (Bio-Rad) with the following settings: 2.5 kV, 200 Ω, 25 μF. Cultures were recovered with LB or SOC media for 1 h at 37°C (30°C for *P. putida*), plated on LB-agar with the proper antibiotic and grown overnight at the required temperature for each host.

#### Pseudomonas aeruginosa

100 ml of *P. aeruginosa* was grown at 37°C until saturation. Cells were centrifuged and washed three times at room temperature. A final washing step was made with 20% Sucrose (Sigma-Aldrich) and the cells were resuspended in 1/100^th^ culture volume of room temperature water. Mixture proportion and electroporation settings were the same used in the coliforms section. Cells were recovered for 1 h at 37°C in LB media cells, plated in LB-gentamycin and grown overnight.

#### Acinetobacter baylyi

*A.baylyi* ADP1-ISx was grown overnight in 3 mL LB medium, at 30°C and 250 rpm in a culture tube. The following day, 70 μL of this culture was added to 1 mL of fresh LB medium in a culture tube and incubated for 3 h along with the recombitron plasmids. 250 ng of plasmid DNA added to 1 mL of culture was sufficient to obtain transformants. After the 3 h, 150 μL of the medium was plated on 25 μg/mL Kanamycin, spread with a cell spreader, and allowed to dry briefly before moving to the incubator.

#### Lactobacillus reuteri

*L. reuteri* was grown in 10 mL MRS for 16 h at 37°C and sub-cultured (Initial OD_600_=0.1) in 40 mL pre-warmed MRS. Cell concentration at OD_600_=0.6 was used for preparing electro-competent cells by following previously established protocol (van Pijkeren and Britton, 2014). Plasmids or LCR ligates were electroporated (2.5 kV, 25 µF, 400 Ohm, and 2 mm gap) into *L. reuteri* electro-competent cell. Following 3 h recovery in 1 mL MRS, cells were plated on MRS-agar containing either 5 μg/mL chloramphenicol for pVPL3017 or 5 μg/mL erythromycin for recombitron-pNZ9530 LCR ligates followed by incubation in the hypoxic chamber (5% CO_2_, 2% O_2_, and N_2_ balance).

#### Mycobacterium smegmatis

*M. smegmatis* mc^2^155 was grown in 300 ml of 7H9 with ADC, CB (50 µg/ml), CHX (10 µg/ml) and Tween 80 (0.05%) at 37°C until OD_600_ was between 0.8 to 1. Cells were incubated on ice overnight. Cells were then centrifuged at 2000 g and washed with ice-cold 10% glycerol with 0.05% Tween 80 at 4°C three times. Finally, cells were resuspended in 1/125^th^ culture volume of ice-cold 10% glycerol with 0.01% Tween 80. 100 μl of the cells were mixed with a 0.15-0.35 μg recombitron plasmid in an ice-cold 0.2-cm gap cuvette (Bio-Rad) followed by electroporation. Electroporation was done using an ECM 630 Electro Cell Manipulator (BTX Harvard Apparatus) with the following settings: 2.5 kV, 1000 Ω, 25 μF. Cells were recovered in 900 μl of 7H9 with ADC and Tween 80 (0.05%) for 3 h at 250 rpm in 37°C. Cells were then plated on 7H10 with ADC and Kanamycin (20 µg/ml) 3 d at 37°C for visible isolated colonies.

#### Cutibacterium acnes

*C. acnes* KPA171202 was transformed as previously described (Knödlseder et al., 2024). In short, *C. acnes* was inoculated to an OD of 0.1 and grown for 24h at 37 degrees, anaerobically, 110 rpm. After 24 hours, cells were treated with 10 ug/mL penicillinG and 0.4 M sucrose and incubated for 4 h more. Then, cells were spun down at 1,700g for 10 min at 4 °C and washed with an equal volume of EPB (272 mM sucrose, sterile-filtered) followed by a second spin under the same conditions. Cells were resuspended in 1 ml of EPB and washed another five times at 9,400g for 1 min at 4 °C. Then, the pellet was resuspended in residual liquid. Competent cells were diluted 1:4 in EPB to a final volume of 50 ul, and 1000 ng of DNA was added to each sample. Cells were transferred to a precooled 0.1-cm electroporation cuvette (Bio-Rad) and electroporated at 1.5 kV, 400 Ω and 25 uF. Cells were recovered in 100 ul of BHI medium and plated on a Brucella agar plate without antibiotic selection. After 24 h of anaerobic incubation at 37 °C, cells were resuspended in 1 ml of BHI to remove remaining non-incorporated DNA, spun down for 5 min at 1,700g and resuspended in 100 ul of BHI. Then, 100 ul of resuspended cells were plated on a Brucella agar plate supplemented with 10 ug ml−1 erythromycin per plate. Transformants were obtained following anaerobic incubation at 37 °C for 7 days.

### Conjugation of plasmids into bacterial species

#### Erwinia amylovora and Pseudomonas syringae

Plasmids were transformed into chemically competent cells of the conjugation donor strain WM3064 (prepared using ZymoResearch “Mix and Go!” Kit). Conjugations were performed by washing donor cells in modified King’s B supplemented with DAP twice, then mixing approximately 1 OD of donor with 1 OD of recipient on 0.45-µm nitrocellulose filters (Millipore) overlaid on modified King’s B agar plates supplemented with DAP and incubated overnight. Filters were collected and mixed with 1 mL modified King’s B liquid, which was diluted and plated with appropriate antibiotics.

#### Shewanella oneidensis, Vibrio natriegens and Aeromonas hydrophila

Recombitron plasmids were first transformed into *E. coli* WM3064 by electrotransformation. 10 ml of *E. coli* WM3064 was grown at 37°C until saturation. Cells were centrifuged and washed three times with 10% glycerol at room temperature and finally resuspended with 250 µl 10% glycerol. 0.1-1 µg of recombitrons plasmids were mixed with 33 µl cell aliquots and transferred to 0.1-cm gap electroporation cuvettes and electroporated immediately at 1.8 kV for 5 ms. Cultures were recovered with 500 µl SOC media for 2 h at 37°C, plated on LB-agar with the proper antibiotic and grown overnight at 37°C. For conjugation, 1 ml cultures of *E. coli* WM3064 harboring different recombitron plasmids and 1 ml of *S. oneidensis* JG2150 were grown at 37°C and 30°C respectively. 250 µl of each WM3064+recombitron culture was centrifuged and washed once with 500 µl LB media. Cell pellets were resuspended with 150 µl JG2150 culture, and the entire volume of cells were plated on LB-agar supplemented with 300 µM diaminopimelic acid and incubated for 8 h at 30°C. A lump of cells were scooped and re-streaked onto LB-agar with the proper antibiotic (50 µg/ml kanamycin for *S. oneidensis* and *A. hydrophila* and 100 µg/ml for *V. natriegens*) for single colonies and grown overnight at 30°C.

### Introduction of plasmids into Streptococcus suis P1/7 by natural transformation

To make *S. suis* P1/7 competent, a nine amino acid peptide (GNWGTWVEE; GenScript) with ≥ 95% purity was used. The peptide was dissolved in DMSO at a final concentration of 5 mM, taking in consideration their specific purity. *S. suis* was grown overnight in THB broth (10g/l Beef Heart, Infusion, 20 g/l Peptic Digest of Animal Tissue, 2g/l Dextrose, 2g/l Sodium Chloride, 0.4 g/l Disodium Phosphate and 2.5 g/l Sodium Carbonate) at 37°C. Next day, a 1∶40 dilution was made into pre-warmed medium, and grown at 37°C without shaking. When OD_600_ reached a value between 0.04-0.06, 100-µl samples were removed from the main culture, and 1-2 µg of recombitrons plasmids were added to the bacteria along with 5 µl of the peptide at a final concentration of 250 µM. After 2 hours of incubation at 37°C the samples were plated in THB or chocolate agar plates with chloramphenicol and grown for 24-48 hours.

### Construction of Klebsiella pneumoniae ΔrecJ/ ΔsbcB strain

Oligonucleotide recombineering was used to construct the *Klebsiella pneumoniae* ΔrecJ/ΔsbcB strain. 70 bp oligonucleotides with phosphorothioate bonds at both ends to protect against native exonucleases were designed to make knockouts in *recJ* and *sbcB* genes (Supplementary Table 6). *K. pneumoniae* cells containing pORTMAGE-Ec1 were grown overnight and next day cultures were diluted 1:100 and grown until OD_600_ ∼ 0.3. At that point, SSAP expression was induced for 1 hour with 1 mM m-toluic acid. Cells were then prepared for transformation as described above and 50 μL of 10 μM of each oligo was added to electrocompetent cells. This mixture was transferred to an electroporation cuvette with a 0.2-cm gap and electroporated immediately using a Gene Pulser (Bio-Rad) with the following settings: 1.8 kV, 200 Ω, 25 μF. Cultures were recovered in LB media for 1 h and then 4 mL of LB with 1.25x fold kanamycin were added for outgrowth at 37°C for 3 hours. Dilutions of the cells were plated on LB-kanamycin and individual colonies were screened to detect the desired mutations. Briefly, 100 µL of 20-30 individual colonies were grown overnight. 25 μl of the culture was mixed with 25 μl of water and incubated at 95°C for 10 min. A volume of 1 μl of this boiled culture was used as a template in 25-μl reactions with primers flanking the edit site. Mutations in *recJ* and *sbcB* genes were verified using Sanger sequencing.

### RT-DNA expression and gel analysis

Production and analysis of the engineered RT-DNA was performed similarly to a previous work (Khan et al., 2024). Briefly, recombitron plasmids were transformed into *E. coli* bMS346, *C. freundii* ATCC 8090, *K. pneumoniae* ATCC 10031 and *P. aeruginosa* PAO1 for expression. An individual colony was used to start a pre-inoculum that was grown overnight in 3 ml LB with the proper antibiotic. A 1:100 dilution of the saturated culture was grown in a flask with 25 ml of LB. The culture was grown for 2h at 37°C to reach OD_600_ between 0.4 and 0.6 and then induced with 1 mM m-toluic acid (Sigma-Aldrich). After 5 h, OD_600_ was measured and the culture was centrifuged at 4,000 rpm for 10 minutes, supernatant was discarded and bacterial pellet were collected for RT-DNA analysis. RT-DNA was recovered using a Qiagen Plasmid Plus Midi Kit and eluted into a volume of 150 μl. Volume of RT-DNA prep was adjusted based on bacterial OD, measured at the point of collection, to normalize the input before loading into Novex TBE–urea gels (15%; Invitrogen). The gels were run (45 min at 200 V) in a preheated (>75 °C C) TBE running buffer. Gels were stained with SYBR Gold (Thermo Fisher Scientific) and then imaged on a Gel Doc Imager (Bio-Rad). To quantify the amount of RT-DNA production relative to retron-Eco1, a retron-Eco1 expressing strain was included in every batch of culture grown, and the resulting prep was always run on the same gel as the experimental retron for quantification. The density of the band produced by each retron was quantified with ImageJ software.

### Retron-mediated recombineering experiments

Once the distinct host cells were transformed with the specific protocol, experiments were conducted in 0.5 ml, deep 96-well plates or in either 3 ml or 10 ml in tubes with the appropriate media for each one the tested bacterial species. For biological replicates, 3 individual colonies of every recombitron were used in this study. Cells were grown until saturation at the required temperature for each organism. Next day, a 1:1000 dilution of the cultures was grown in the presence of the inducer (if required): 1mM of m-toluic acid (Sigma-Aldrich) for plasmids regulated by the XylS/Pm system; 1 mM IPTG (GoldBio) for plasmids regulated by the lacI/P_lac_ system; 50 mM sucrose (Sigma-Aldrich) and 20 ng/ml pSIP induction peptide (MAGNSSNFIHKIKQIFTHR from Peptide 2.0 Inc) in mMRS (modified MRS without sugar) for inducing P_suc_ and P_orfx_ in *L. reuteri*, respectively. Cultures were grown overnight for the specific time that a host requires to reach saturation.

For genotyping and retron-mediated recombineering efficiency for *L. reuteri*, recombinant colonies grown on MRS-agar containing 25 µg/ml rifampicin in the hypoxic chamber (24 h at 37°C) were screened by colony-PCR using mismatch amplification mutation assay (MAMA)-PCR oligos (oVPL304, 305, and 306) (van Pijkeren and Britton, 2012). The ratio of rifampicin-resistant colonies confirmed by MAMA-PCR per total viable cells was used to calculate % recombineering efficiency.

For *M. smegmatis*, in a 50 ml baffled flask, 8 μl of a recombitron plasmid-carrying saturated culture was used to inoculate 8 ml of 7H9 with succinate (0.2%), acetamide (0.2%), Kanamycin (20 µg/ml), and Tween80 (0.05%). The inoculated medium was incubated at 37°C and 250 rpm. At 24 h, 48 h, and 72 h, a 1-1.5 ml sample was collected and OD600 measured. For the samples with low OD600, they were concentrated by centrifugation and resuspended in much smaller volumes to increase cell density. The subsequent steps are described in the above section starting from the part of a volume of 25 μl of culture.

For *C. acnes*, after 7 days of anaerobic incubation at 37 degrees, colony PCR of transformants was performed to verify presence of plasmid DNA and re-streaked on a new Brucella agar plate supplemented with 10 ug/mL erythromycin. After 48 h, transformants were inoculated in 10 mL BHI media in a 25 cm2 flask supplemented with erythromycin to an initial OD600 of 0,1 and grown for 24h, 37 degrees, anaerobically, 110 rpm. A minimum of three passages was performed per colony by re-inoculating every 24h to OD600 of 0,1. Samples were taken after every passage and treated as described above.

### Sequencing and editing rate quantification

For sequencing and editing quantification for all hosts used in this study, except *L. reuteri*, a volume of 25 μl of culture was collected just after the experiment that was mixed with 25 μl of water and incubated at 95°C for 10 min. A volume of 1 μl of this boiled culture was used as a template in 25-μl reactions with primers flanking the edit site, which additionally contained adapters for Illumina sequencing preparation. These amplicons were indexed using primers listed in Supplementary Table 6 and sequenced on an Illumina NextSeq instrument and processed with custom pipeline to quantify the percentage of precisely edited genomes.

## ACKNOWLEDGEMENTS

We thank the lab of Joseph Bondy-Denomy for providing the *Pseudomonas aeruginosa* strain PAO1. We thank Professor Marcelo Gottschalt and Sonia Lacouture at University of Montreal for providing the *Streptococcus suis* strain P1/7. We also thank the lab of Jerry M. Wells at Wageningen University for supplying pSStarget vector. Work was supported by funding from the National Science Foundation (MCB 2137692), the Gary and Eileen Morgenthaler Fund, the Gordon and Betty Moore Foundation, and the Robert J Kleberg, Jr. and Helen C. Kleberg Foundation. S.L.S. is a Chan Zuckerberg Biohub – San Francisco Investigator. Contributions made by the Mutalik lab were made possible by the Biopreparedness Research Virtual Environment (BRaVE) Phage Foundry at Lawrence Berkeley National Laboratory is based upon work supported by the U.S. Department of Energy, Office of Science, Office of Biological & Environmental Research under contract number DE-AC02-05CH11231 Contributions made by the van Pijkeren lab were made possible by grant number R01AT011202 from the National Center for Complementary and Integrative Health (NCCIH) at the National Institutes of Health. Its contents are solely the responsibility of the authors and do not necessarily represent the official views of NCCIH.

## AUTHOR CONTRIBUTIONS

A.G.-D. and S.L.S. conceived the study, design the experimental procedure and centrally coordinated the collaboration with the other labs. A.G.-D. cloned most plasmids used in this study, other than plasmids used in *L. reuteri* experiments, which were cloned by J.-H.O, some plasmids used in S. oneidensis, which were clone by Z.H., and those used in *S. suis* experiments which were clone by L.B.L.. A.G.-D. performed all experiments with *E. coli*, *C. freundii*, *K. pneumoniae*, and *P. aeruginosa*. M.S.J. performed all experiments with *E. amylovora* and *P. syringae*. M.C.W. performed all experiments with *V. natriegens* and *A. hydrophila* and the experiment targeting *flgH* gene in *S. oneidensis*. Z.Y. performed most experiments in *S. oneidensis* with some preliminary results obtained by Y.L. A.G.-D. and Z.Y. performed all experiments in *P. putida*. H.S. performed all experiments in *A. baylyi*. J.-H.O. performed all experiment in *L. reuteri*. L.B.L. performed all the experiments in *S. suis*. C.-C.K. performed all the experiments in *M. Smegmatis*. N.K. performed all experiments in *C. acnes*. V.A., J.A.G., M.G., G.F.H., B.K.K., B.K., V.K.M., J.P.P., and S.L.S. supervised the experiments in the distinct bacterial species. A.G.D. prepped samples for NGS and ran the sequencing libraries. A.G.-D. analyzed the data. A.G.-D. and S.L.S. wrote the manuscript with input from all authors.

## COMPETING INTERESTS

The authors declare no competing interests.

## DATA AND CODE AVAILABILITY

Sequencing data associated with this study are available in the NCBI SRA (Bioproject PRJNA1269658) http://www.ncbi.nlm.nih.gov/bioproject/1269658

## SUPPLEMENTARY FIGURES

**Supplementary Figure 1.**
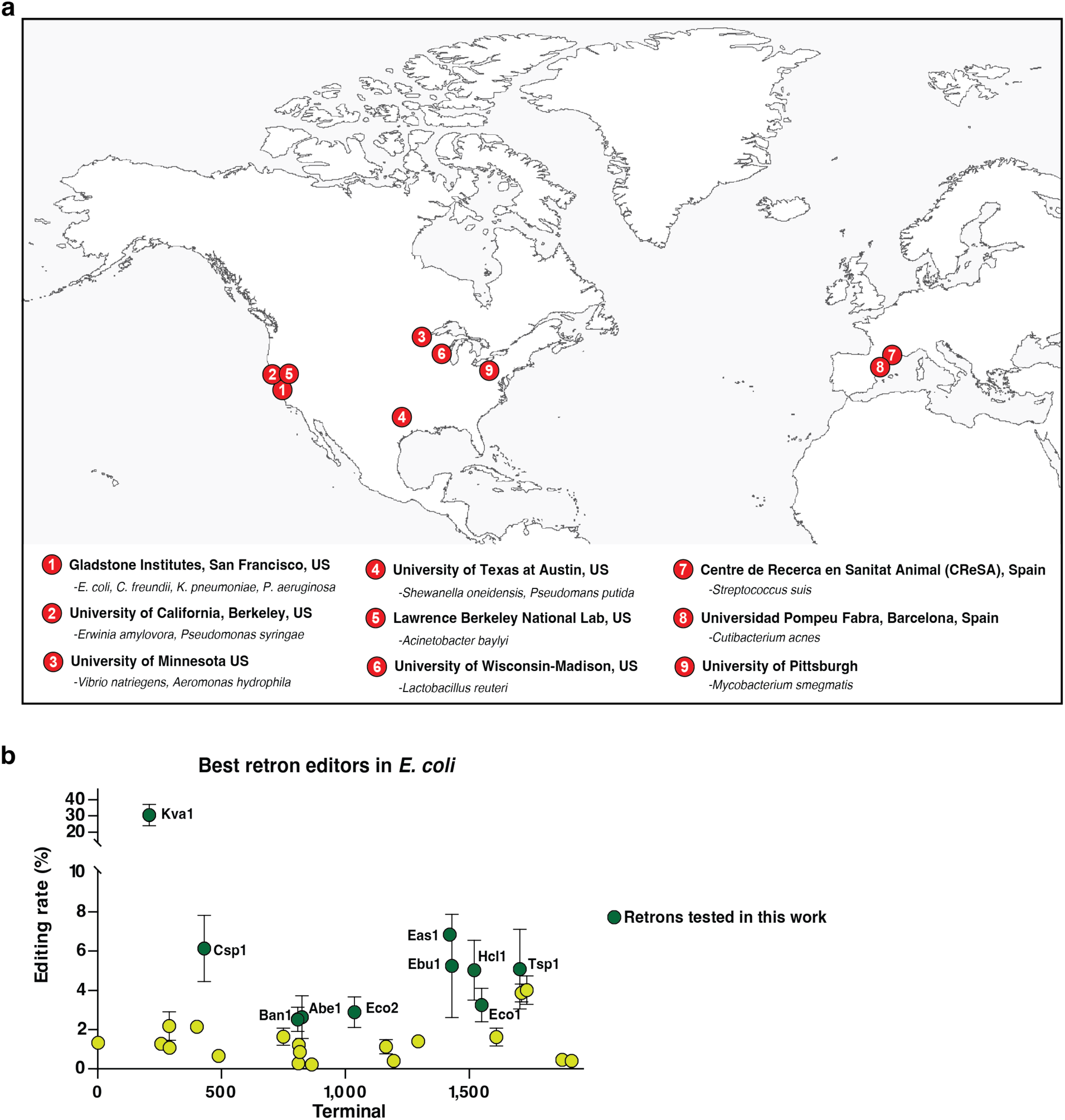
Geographic distribution of the research lab involved in the recombitron portability project and previous performance of the retrons used in this survey. **a**, Map showing the Academic Institutions involved in this project and the species assayed in each one of the research labs. **b**, Precise editing rate across retrons for bacterial genome recombineering. Points show mean ± standard error of the mean for three biological replicates. The name of the 10 selected retrons is indicated. Adapted from Khan et al., 2024.

**Supplementary Figure 2.**
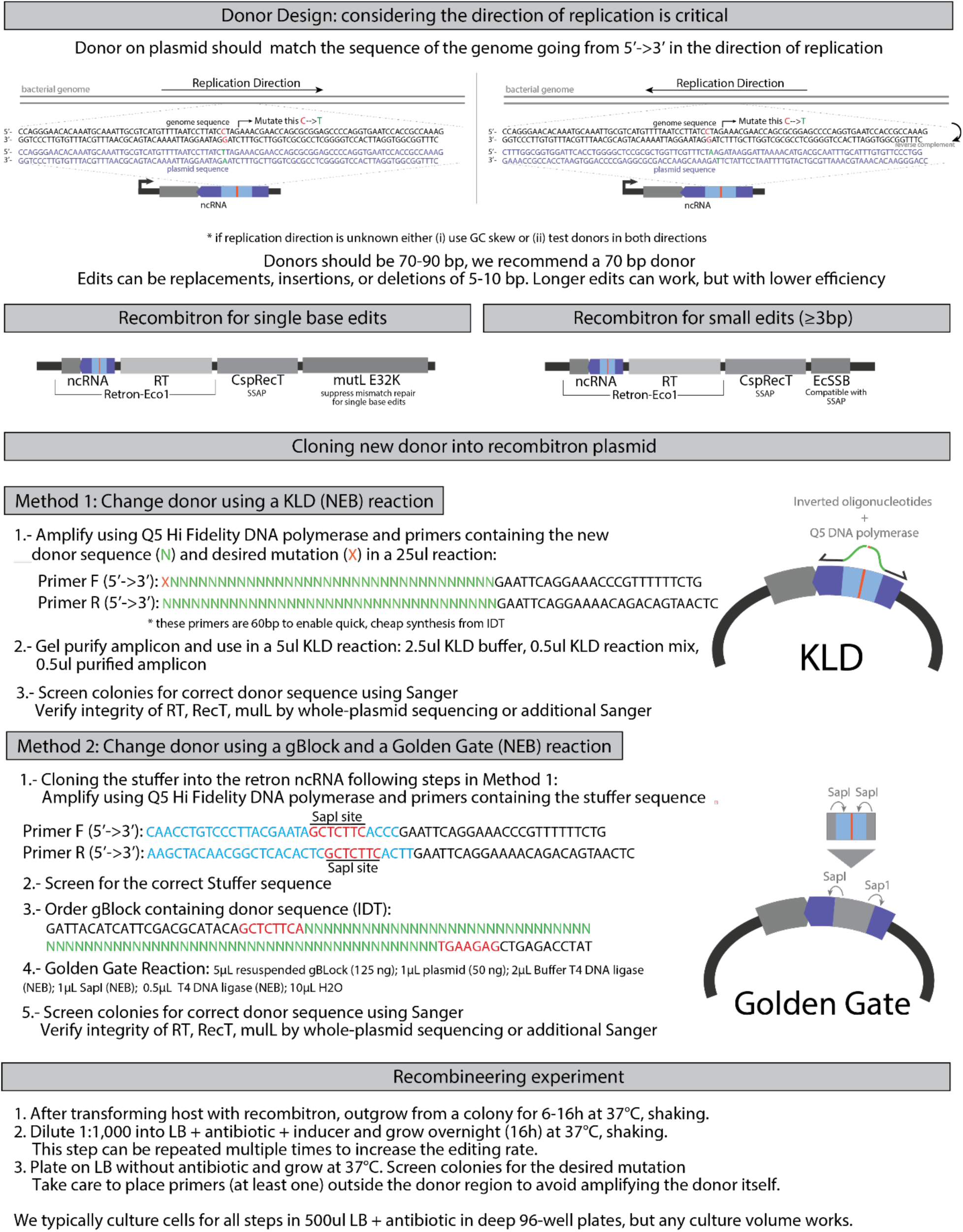
**Protocol to design, construct and assay a recombitron for genome editing.**

**Supplementary Figure 3.**
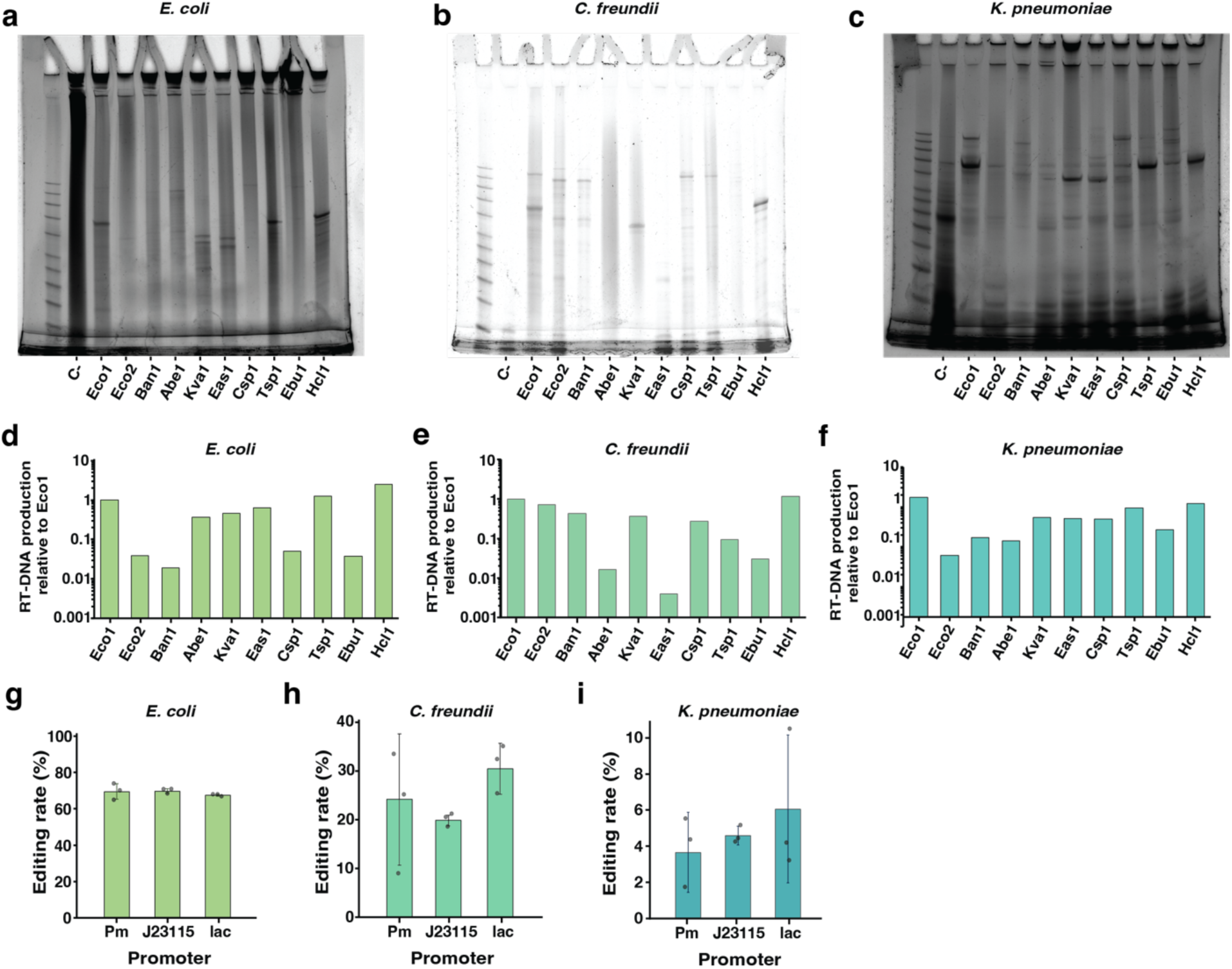
Retron-mediated recombineering in coliforms. **a**, Uncropped gel from **Fig 2b**, showing RT-DNA production from the set of 10 recombitrons in *E. coli*. **b**, Uncropped gel from **Fig 2c**, showing RT-DNA production from the set of 10 recombitrons in *C. freundii*. **c**, Uncropped gel from **Fig 2d**, showing RT-DNA production from the set of 10 recombitrons in *K. pneumoniae*. **d**, Quantification of RT-DNA production by density, relative to retron-Eco1 in *E. coli*. **e**, Quantification of RT-DNA production by density, relative to retron-Eco1 in *C. freundii.* **f**, Quantification of RT-DNA production by density, relative to retron-Eco1 in *K. pneumoniae*. In **d-f**, Density of the band produced by each retron was quantified with ImageJ software. **g**, Quantification of precise genome editing to make a 5 bp deletion in the *lacZ* gene in *E. coli* using Pm, lac or J23115 promoter. **h**, Quantification of precise genome editing to make a 5 bp deletion in the *lacZ* gene in *C. freundii* using Pm, lac or J23115 promoter. **i**, Quantification of precise genome editing to make a 5 bp deletion in the *lacZ* gene in *K. pneumoniae* using Pm, lac or J23115 promoter. In **g-i**, data were quantified by sequencing after 24 h of editing using Illumina NextSeq, circles show each of the three biological replicates, and errors bars are mean ± standard deviation.

**Supplementary Figure 4.**
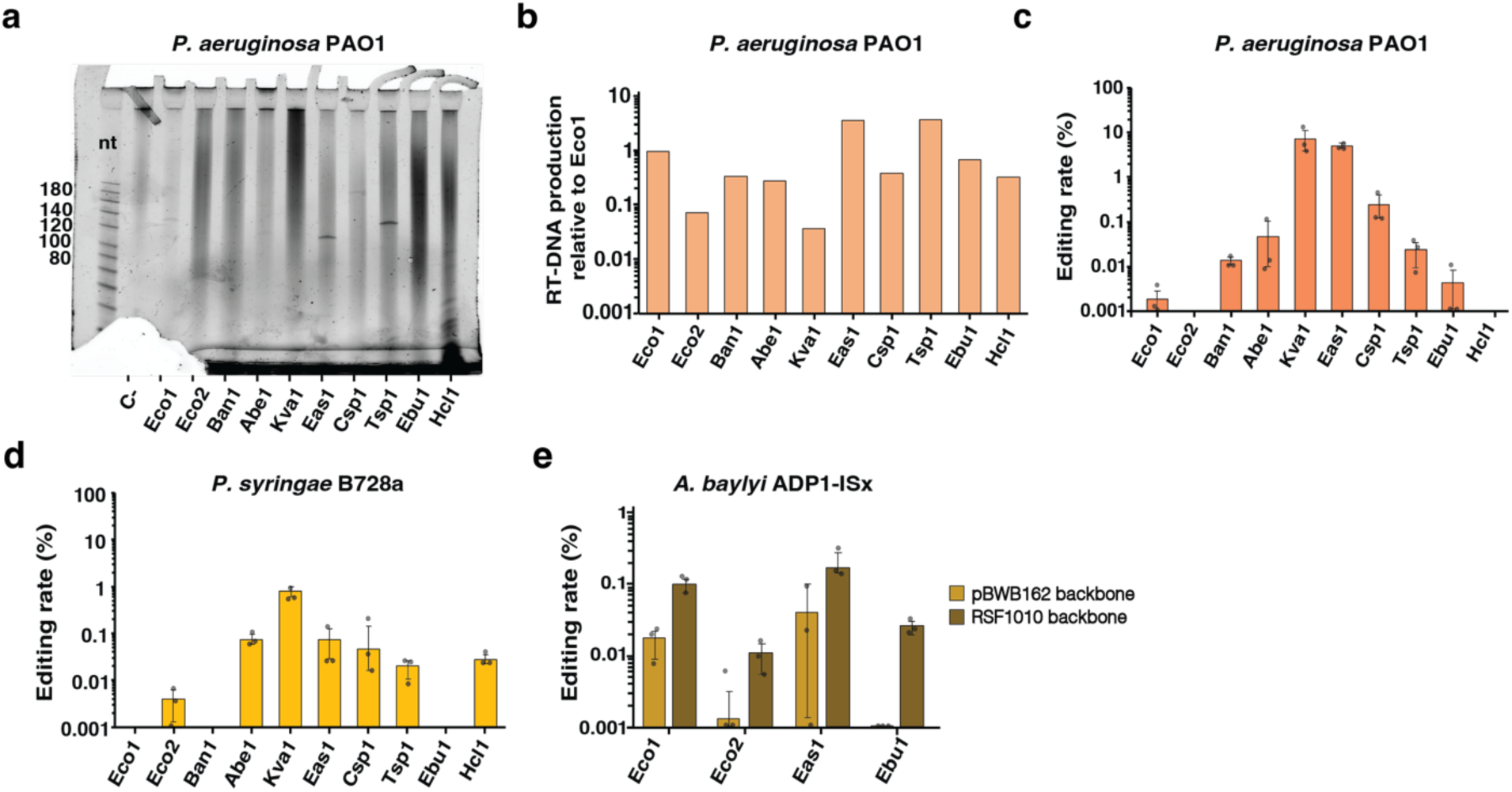
Retron-mediated recombineering in *Pseudomonadales*. **a**, PAGE analysis of RT-DNA production in *P. aeruginosa* PAO1. **b**, Quantification of RT-DNA production by density, relative to retron-Eco1 in *P. aeruginosa.* Density of the band produced by each retron was quantified with ImageJ software. **c**, Quantification of precise genome editing to make a 4 bp deletion targeting the *phzM* gene in *P. aeruginosa* PAO1 using the 10 recombitrons set with pBBR1 inverted origin of replication. **d**, Quantification of precise genome editing to make a 5 bp deletion in a non-essential intergenic region in *P. syringae* B728a using the 10 recombitrons set with pBBR1 inverted origin of replication. **e**, Comparison of precise genome editing to make a 4 bp deletion targeting a glycosyltransferase gene in *A. baylyi* sFAB6437 using recombitrons expressed from pWBW162 or RSF1010 backbone. In **c-e**, data were quantified by sequencing after 24 hours of editing using Illumina NextSeq, circles show each of the three biological replicates, and errors bars are mean ± standard deviation. Additional statistical details are presented in Supplementary Table 3.

**Supplementary Figure 5.**
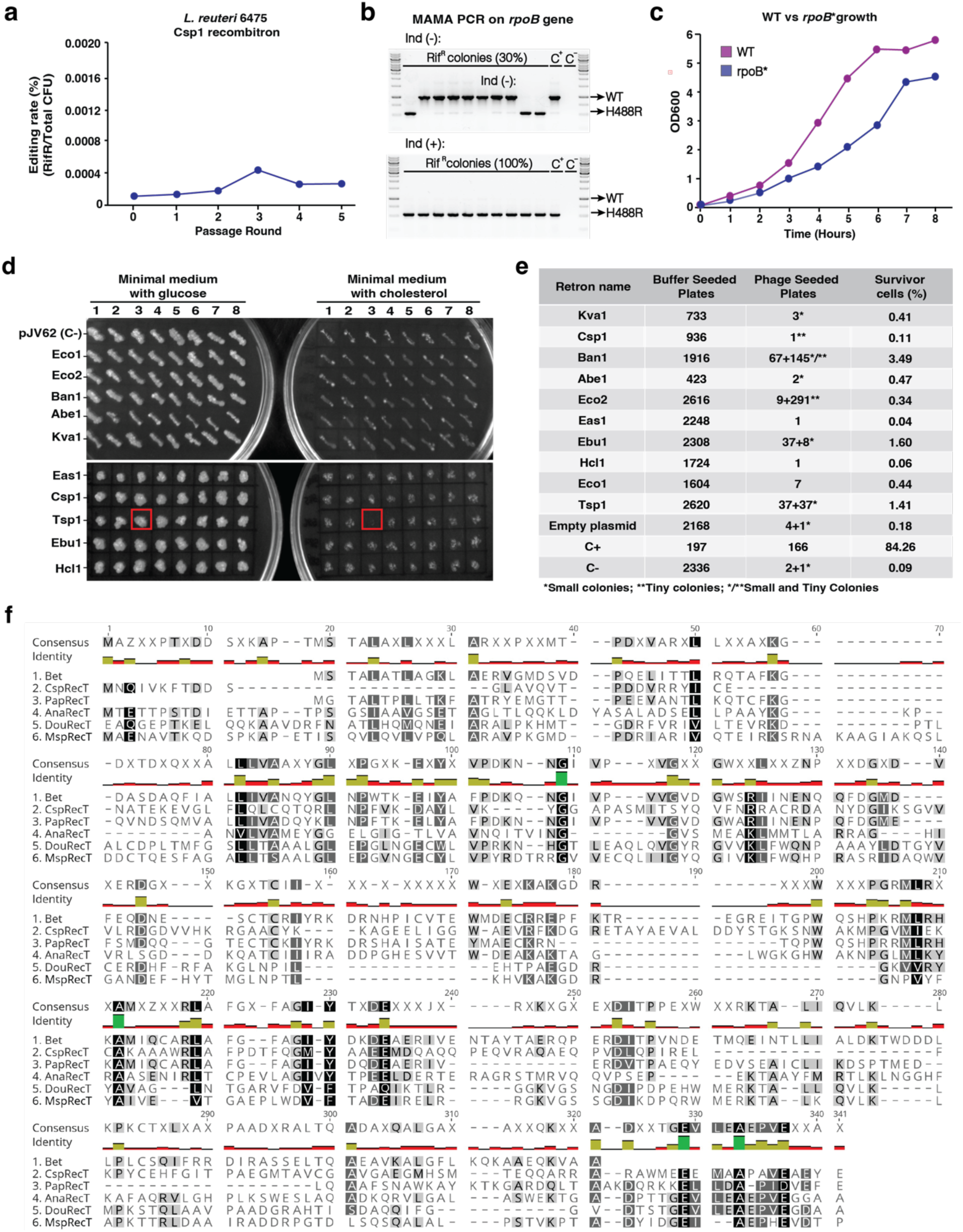
Retron-mediated recombineering in *Bacillota* and *Actinomycetota*. **a**, Quantification of precise genome editing to make a single-point mutation in *rpoB* (H488R) gene in *L. reuteri* 6475 using a single plasmid assay with-Csp1 recombitron. The experiment was performed for 6 passage rounds in mMRS with P_suc_ and P_orfx_ inducers. Editing rates were calculated as % of rifampicin resistant colonies respect to total number of colonies in no-antibiotic plates **b**, Mismatch amplification mutation assay (MAMA) PCR to confirm targeted mutations in the *rpoB* gene. Rifampicin-resistant colonies were used as template for the PCR. Top: screen of 10 colonies with no recombitron induction. Bottom: screen of colonies with recombitron induction. Wild type (WT) and mutant (H488R) bands are indicated with an arrow. **c**, Time course of the OD_600_ of wild type versus mutated *rpoB* gene (H488R) cells. **d**, Minimal Media (MM) plates with glucose or cholesterol with 8 individual colonies of each one of the 10 recombitrons and a negative control (pJV62). No growth of colony 3 on MM with cholesterol indicates that cell was edited by-Tsp1 recombitron. **e**, Quantification of genome editing by infecting with Mycobacteriophage Brilliant. Edited cells are resistant to phage infection. The number of total colonies in the presence of buffer or the phage is indicated. The editing is represented by the percentage of survivor colonies in the phage-seeded plates respect to total colonies in the buffer-seeded plates. **f**, MAFFT Sequence alignment of 6 phylogenetically distinct RecT proteins, including AnaRecT and DouRecT used in *C. acnes* experiments. Sequence shading shows residue conservation and similarity: white text on black background, 80% conserved; grey background, 60% conserved. The alignment was performed using Geneious Prime software.

## Notes

### Competing Interest Statement

The authors have declared no competing interest.

### Summary of Updates

Names, affiliations and funding source was updated

